# Blocking D2/D3 dopamine receptors increases volatility of beliefs when we learn to trust others

**DOI:** 10.1101/2022.06.21.496956

**Authors:** Nace Mikus, Christoph Eisenegger, Chris Mathys, Luke Clark, Ulrich Müller, Trevor W. Robbins, Claus Lamm, Michael Naef

## Abstract

The ability to flexibly adjust beliefs about other people is crucial for human social functioning. Dopamine has been proposed to regulate the precision of beliefs, but direct behavioural evidence of this is lacking. We investigated how a relatively high dose of the selective D2/D3 dopamine receptor antagonist sulpiride impacts learning about other people’s prosocial attitudes in a repeated trust game. Using a Bayesian model of belief updating, we show that sulpiride increased the volatility of beliefs, which led to higher precision-weights on prediction errors. This effect was entirely driven by participants with genetically conferring higher dopamine availability (Taq1a polymorphism). Higher precision weights were reflected in higher reciprocal behaviour in the repeated trust game but not in single-round trust games. This finding suggests that antipsychotic medication might acutely reduce rigidity of pathological beliefs.

## Introduction

Knowing whom to trust with our money, information, or health is central to our personal well-being (Meyer-Lindenberg & Tost, 2012). The ability to form beliefs about other persons’ attitudes from their actions is therefore pivotal for successfully navigating our social world. Inflexible beliefs, particularly about intentions of others, can lead to thoughts of persecution or even paranoid delusions - a hallmark symptom of psychotic disorders (Diaconescu, Wellstein, Kasper, Mathys, & Stephan, 2020; Gromann et al., 2013; Wellstein et al., 2020). Understanding the neurocomputational substrates of social inference is therefore essential for informing pharmacological treatments of psychotic symptoms.

When learning whether to trust another person, we often do so by observing their behaviour across repeated interactions. How behaviours of others affect our overall beliefs about their trustworthiness largely depends on how certain we are about the attitudes that presumably drive others’ actions (FeldmanHall & Shenhav, 2019). For instance, if we firmly believe someone is hostile towards us, a positive gesture coming from them will not much change our belief about them. On the other hand, that same gesture from someone whose intentions we are unsure of, will likely strongly shift what we think about them. This process of belief updating under uncertainty has been formalized within the Bayesian Inference framework, where beliefs are represented as probability distributions and the degree to which new information affects the updating of beliefs is modulated by the precision (the inverse of uncertainty) of those beliefs (Mathys, Daunizeau, Friston, & Stephan, 2011). As in similar computational frameworks (Sutton & Barto, 2017), the belief update is proportional to the deviation of the prediction from the actual outcome, termed as a prediction error (PE), weighted by the precision of prior beliefs. When prior beliefs are highly uncertain, the weight on the PE will be high, and conversely, highly precise prior beliefs lead to a down-regulation of the influence of PE on learning. Inflexibility in changing our beliefs about others can therefore result from high precision of prior beliefs about others’ attitudes. Yet, the neurocomputational and neurochemical mechanisms of regulating uncertainty of beliefs are poorly understood. In this study we examined the effects of the commonly used antipsychotic drug sulpiride, a D2/D3 dopamine receptor antagonist, on the uncertainty of beliefs about another person’s trustworthiness.

Seminal studies in animals have established that mesolimbic dopaminergic circuits carry PE signals that drive belief updating in various contexts (Matsumoto & Hikosaka, 2009; Montague, Dayan, & Sejnowski, 1996; Schultz, 1998). However, dopaminergic midbrain neurons have been shown to be involved in various probabilistic computations that go well beyond phasic signalling of surprising rewarding events. Dopamine responses scale with outcome variance (Schultz et al., 2008; Tobler, Fiorillo, & Schultz, 2005) and reflect temporal and perceptual precision (De Lafuente & Romo, 2011; Fiorillo, Newsome, & Schultz, 2008; Fiorillo, Tobler, & Schultz, 2005). Several computational accounts of brain function suggest that uncertainty or precision coding is the main unifying feature of dopamine’s involvement in belief updating (Friston, Stephan, Montague, & Dolan, 2014; Gershman, 2018; Gershman & Uchida, 2019; Mikhael & Bogacz, 2016). Through encoding of uncertainty of beliefs about the world and what action to perform, dopamine receptors are thought to adjust the weights on PEs and control action selection (Adams, Stephan, Brown, Frith, & Friston, 2013; Babayan, Uchida, & Gershman, 2018). But while there is evidence for the involvement of dopamine receptors in processing uncertainty in action selection (Adams et al., 2020; Eisenegger et al., 2014, 2013; Schwartenbeck, FitzGerald, Mathys, Dolan, & Friston, 2015), no study in humans has yet demonstrated their causal role in regulating the uncertainty of social beliefs and adjusting weights on PEs.

Dopamine receptors within the corticostriatal circuitry are ideally positioned to regulate the PE-related signal propagation and encode precision (Friston, 2008; Yao, Spealman, & Zhang, 2008). D1 and D2 type dopamine receptors in the striatum are believed to be involved in complementary aspects of PE signal propagation (Yao et al., 2008). Whereas D1-type receptors mostly respond to phasic dopaminergic bursts and amplify striatal output, the D2 receptor class (including D3 receptors) plays a gating role in signal propagation (Frank, 2005) by attenuating phasic dopamine release (Ford, 2014; Grace, 2016) and thus by regulating corticostriatal excitation (O’Donnell & Grace, 1994; Yin & Lovinger, 2006). Blocking D2 receptors could bias the activity of the cortico-striatal loops towards the D1 receptor driven pathway and therefore deregulate PE signal propagation, leading to reduced precision of prior beliefs (Adams, Vincent, Benrimoh, Friston, & Parr, 2021).

Although some studies indeed showed that blocking D2-type receptors enhanced learning from positive feedback (Frank & O’Reilly, 2006), led to pronounced PE-related activity in the striatum (Iglesias et al., 2021), and enhanced performance (Eyny & Horvitz, 2003; Jocham, Klein, & Ullsperger, 2011), there is also evidence for attenuated PE coding and greater variability in choice selection (Eisenegger et al., 2014; Jocham, Klein, & Ullsperger, 2014; Pessiglione, Seymour, Flandin, Dolan, & Frith, 2006). The inconsistencies of these findings raise several important considerations. First, when alternative choices are available, it is often unclear whether increased switching between available choice options arises from drug effects on belief updating *per se* or from the effects on decision-making strategies (see for instance (Zhang, Lengersdorff, Mikus, Gläscher, & Lamm, 2020)). Second, D2 dopamine receptors have a higher affinity for dopamine (Richfield, Penney, & Young, 1989) and doses of D2 antagonists commonly used in studies with healthy volunteers might not be enough to sufficiently block the D2 receptor driven regulation of the PE signal (Bressan et al., 2003). Third, administration of compounds binding to dopamine receptors can have different and even opposing effects on learning and decision-making, depending on genetic variation in baseline dopamine function (Cohen, Krohn-Grimberghe, Elger, & Weber, 2007; Cools & D’Esposito, 2011; Eisenegger et al., 2013). And finally, beyond the methodological limitations of previous work, most studies with dopamine receptor antagonists have looked at learning about abstract stimulus-outcome associations using secondary rewards, which makes the translation to more complex social interactions questionable.

In light of these considerations, the present study administered a relatively high dose of the selective D2/D3 receptor antagonist sulpiride (800 mg) or placebo in a randomized double-blind parallel group design to 78 male participants, preselected based on their Taq1a polymorphism. The drug dose was chosen to maximise the blockade of postsynaptic dopamine D2 receptors while still being safe (Takano et al., 2006). Most previous work used doses of 400 mg which leads to an occupation of approximately 30% of D2 receptors (Mehta, Montgomery, Kitamura, & Grasby, 2008). Using 800 mg leads to more than 60% occupancy and increases the likelihood of sufficiently blocking the effect of D2 receptors. Furthermore, as mentioned above, the effect of D2 antagonists often interacts with baseline variation on dopamine function (Cohen et al., 2007; Westbrook et al., 2020). Taq1a polymorphism is one of the most widely investigated genetic variations of the D2 receptor. Individuals with at least one A1 minor allele have been shown to have higher presynaptic dopamine availability (Laakso et al., 2005) and reduced D2 receptor density in some subdivisions of the striatum (Gluskin & Mickey, 2016; Smith et al., 2017). Blocking D2 receptors might therefore have a stronger effect on belief updating in that genetic subgroup.

We investigated social learning by asking the participants to learn about other players’ trustworthiness through a repeated trust game (Fig 1a). In the trust game the investor may choose to transfer any portion of their monetary endowment to the trustee (Berg, Dickhaut, & McCabe, 1995).

**Fig. 1.**
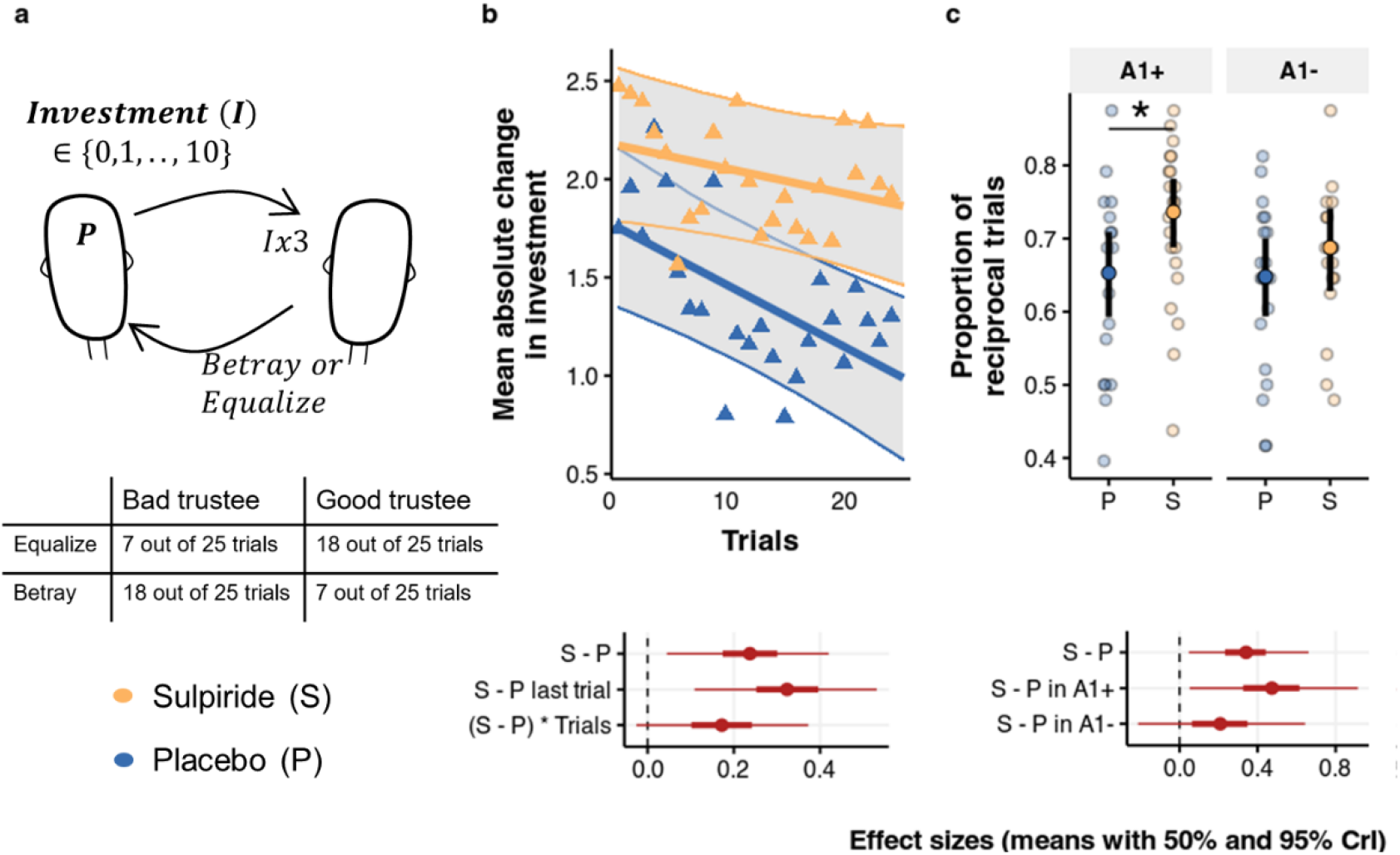
Effects of sulpiride on behaviour in the Repeated Trust Game. **a**, The participants played 25 trials with two trustees. Each trial started with an endowment of 10 points to both players. On each trial they could invest any integer between 0 and 10. The trustee received a tripled amount of the investment and could decide to either equalize payoff or betray the other player and keep all the points for himself. The trustees were pre-programmed to be either “good” or “bad”. **b**, Mean and 95% CrI of absolute change of investment from one trial to the next for both treatment groups based on a Bayesian multilevel model, plotted over raw means for each participant (Δ). Corresponding effect sizes show a main effect of sulpiride, a larger effect of sulpiride on the last trial and a marginal effect of sulpiride on the slope across trials. **c**, Mean and 95% CrI of reciprocal trials (defined as trials where investment was increased following positive feedback, or decreased following negative feedback) based on a Bayesian logistic multilevel model, plotted over raw proportion of reciprocal trials for each participant. Effect sizes in logodds, show a main effect of the drug, driven by the A1+ group.

The transferred points are then multiplied by the experimenter before being passed on to the trustee. The trustee can then either reciprocate in a way that equalizes the payoff of the two players or betray and keep everything. Participants in our study played 25 rounds of the trust game as investors against two other players that were preprogramed to mostly equalize (“good trustee”) or mostly betray (“bad trustee”). Importantly, we told the participants that the other players had given their answers weeks prior to the study day and therefore their decision to equalize or betray did not depend on the participant’s investment. With this procedure we increased the likelihood that their investments reflected the degree of uncertainty they had about the other player’s response and were not confounded by strategic investment strategies, or exploratory action policies.

The main goal of the study was to test the assertion that blocking D2-type receptors increases belief updates by reducing the precision of beliefs about others, whereby we hypothesized that this effect will be more pronounced in participants with genetically conferring higher endogenous dopamine levels. The results section of the paper is structured as follows: we first looked at how sulpiride affects investment updates and how this effect is moderated by the Taq1a genotype. We then examined how the updates relate to the back-transfer of the trustee, by looking at the effects of the drug and genotype on reciprocal behaviour. We then turn to computational modelling to determine how sulpiride affects the course of each participant’s uncertainty around the other player’s trustworthiness. Finally, to control for effects of our drug manipulation on sensitivity to social feedback unrelated to learning, we surveyed data from two single-round social interaction tasks, targeting positive and negative reciprocal behaviour.

## Results

### D2/D3 receptor antagonism increases investment updates

We employed a Bayesian multi-level linear model predicting absolute change in investment from the previous trial, including variables for Treatment (sulpiride or placebo), Trial and their interaction as predictors (refer to supplementary material for outcomes of alternative models). Fig 1b shows that, following sulpiride administration, participants on average updated their investments more than participants in the placebo group (b = 0.633, 95% Credibility Interval (CrI) [0.117, 1.115], proportion of the posterior distribution of the regression coefficient below 0 being P(b<0) = 0.005), with an effect size d = 0.239 (95% CrI [0.045, 0.42]). The difference in investment updates was most apparent in the last trial of the task (b = 0.863, 95% CrI [0.289, 1.411], P(b<0) = 0.002, d = 0.325, 95% CrI [0.109, 0.531]) and we also found a small effect size on the Trial*Treatment interaction (b = 0.457, 95% CrI [-0.069, 0.99], P(b<0) = 0.047, d = 0.172, 95% CrI [-0.026, 0.373]). As participants learned about the trustees, changes of investments from one to the next trial reduced, and this decrease across time was less pronounced in the sulpiride group.

To examine whether the effects of the drug were moderated by the Taq1a polymorphism we ran another model including a variable for Taq1a-specific genotype and Trustee as predictors with the four-way interaction between the two new variables, Treatment and Trial, including a random intercept and slope for the Trustee (Supplementary Fig. 1a, Supplementary Table 4). Again, we found a main effect of treatment (b = 0.595, 95% CrI [0.112, 1.098], P(b<0) = 0.008), and a significant three-way interaction between Treatment, Genotype and Trial number (b = 0.053, 95% CrI [0.01, 0.098], P(b<0) = 0.007), while the two-way interaction Treatment x Genotype was not significant (b = -0.284, 95% CrI [-1.266, 0.708], P(b>0) = 0.287). These analyses suggest that on average sulpiride affected investment updates comparably across both genotype groups, but in contrast to the A2 homozygotes, the effect in the A1+ group was time dependent.

### D2/D3 receptor antagonism increases sensitivity to social feedback in the A1+ group

To further understand how investment updates related to back-transfer from the trustee, we defined reciprocal trials as trials where participants either increased investments (or repeated the maximal investment of 10 points) following positive feedback and decreased investments (or repeated an investment of 0 points) following a betrayal (Fig. 1c, for exact definition see Supplementary Note 2). We found that sulpiride led to a higher proportion of reciprocal trials (*b*_*logodds*_= 0.339, 95% CrI [0.048, 0.661], P(*b*_*logodds*_ <0) = 0.012), and this effect was significant in the A1+ group (*b*_*logodds*_= 0.469, 95% CrI [0.052, 0.914], P(*b*_*logodds*_<0) = 0.015) but not in the A1-group (*b*_*logodds*_= 0.209, 95% CrI [-0.212, 0.643], P(*b*_*logodds*_<0) = 0.162); however, the interaction between the drug and genotype was not significant (*b*_*logodds*_ = -0.263, 95% CrI [-0.867, 0.329], P(*b*_*logodds*_>0) = 0.186). Furthermore, we found some support for a dose response effect, whereby sulpiride serum levels in the blood correlated with reciprocal trials in the A1+ group (b = 0.185, 95% CrI [-0.04, 0.41], P(r<0) = 0.05), but not in the A1-group (Supplementary Table 1). Similar, albeit weaker, effects were found when we examined to what extent the signed investment update was dependent on the back-transfer and how this differed across the drug and genotype groups (Supplementary Fig. 1b). Note that this higher change of investments from one trial to the next did not lead to different investments on average across the four drug/genotype groups across the entire duration of the task (Supplementary Fig 2).

This suggests that sulpiride increases sensitivity to social feedback when learning about others. To determine whether and how this how this behavioural pattern relates to the uncertainty of participants’ beliefs about the other persons’ trustworthiness, we explicitly modelled the participants’ trial by trial evolution of beliefs with a Bayesian belief model. In our modelling approach, we considered that a similar behavioural pattern could also have emerged from increased uncertainty around investment selection, or simply have been due to the degree to which the beliefs about the other player were used to guide investment selection.

### Computational framework

The belief model generates a trial-wise sequence of our participants’ beliefs about the trustworthiness of two trustees as well as the uncertainty (or precision) surrounding those beliefs (Fig 2, (Mathys et al., 2011, 2014)). We estimated a participant-specific parameter *ω*, called *belief volatility*, that describes how each participant’s precision of beliefs evolved across time and consequently determined the relative rigidity (or flexibility) of beliefs. Higher belief volatility *ω* implies higher uncertainty (lower precision) of prior beliefs, meaning higher learning rates and stronger shifts in beliefs throughout the task (see two example participants with different *ω* values in Fig. 2b). The belief model was fitted to the data through an ordinal logistic response model with two additional participant-level parameters. The choice uncertainty parameter *γ* is a probability weight that determines how feedback probability maps on the investment. Higher values imply an investment distribution centred around extremes (i.e., investing 0 and 10) and lower values imply a more dispersed investment distribution and more uncertainty or stochasticity in action selection. It thus mirrors the inverse temperature parameter in the softmax equation often used in non-ordinal (e.g., binary) choice tasks. The trustworthiness slope (*η*) determines to what degree inferred trustworthiness correlates with investments. Crucially, the computational parameters of the model represent distinct behavioural patterns and can be recovered reliably (Fig. 2c). To determine how noisy trials are represented in the model, we defined mistake trials as trials where participants either decreased their investment after a positive back-transfer or increased their investment after a betrayal (for exact definition see Supplementary Note 1). Importantly, we observed that belief volatility *ω* correlates with reciprocity (r = 0.277, t = 2.476, df = 74, p = 0.016, Fig. 2d) and that the log transformed choice uncertainty parameter *γ* correlates negatively with the proportion of mistake trials (r = -0.592, t = -6.3254, df = 74, p < 10e-3, Fig. 2d) implying higher randomness in investment selection. We also predicted data from the posterior distributions of parameters and confirmed that the model captures the crucial aspects of behaviour (Supplementary Fig. 3a,b) as well as compared the model to an HGF model without the *γ* parameter and a Rescorla-Wagner model and found that it outperforms both models (Supplementary Fig. 3c).

**Fig. 2.**
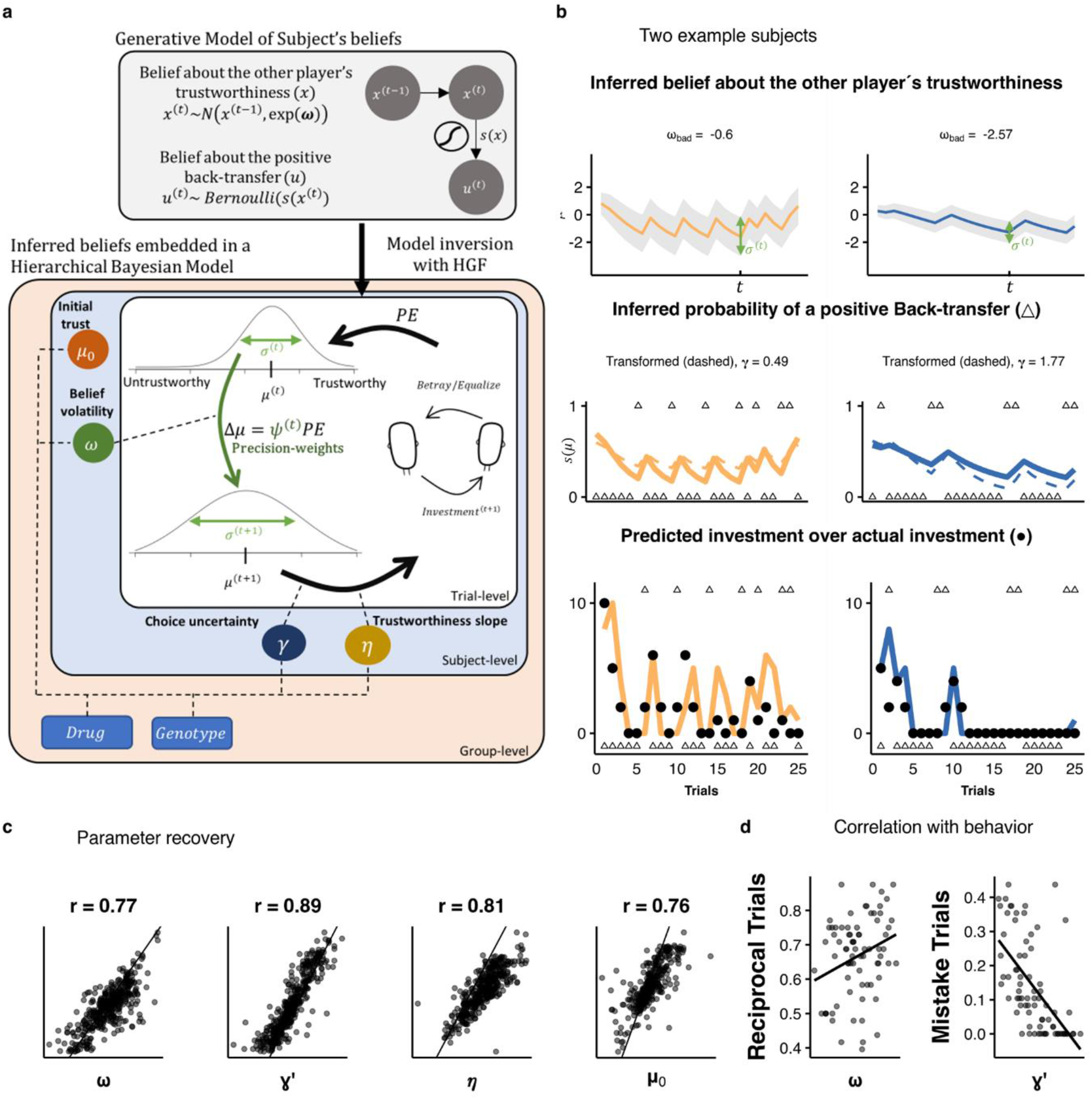
Computational modelling. **a**, We defined a generative model that describes the evolution of participants’ beliefs about the other person’s trustworthiness as Gaussian random walks. We then invert the model using the Hierarchical Gaussian Filter (HGF) and combined the inverted model with an ordinal logistic response model to obtain the likelihood function. The HGF generates trial-level estimations of participants’ beliefs about the trustworthiness of others as Gaussian variables with mean *μ*^(*t*)^ and standard deviation *σ*^(*t*)^. The precision-weights *ψ*^(*t*)^ on the prediction error in each trial are determined by the precision (inverse variance) of beliefs about the other player’s trustworthiness. The evolution of the variances is in turn determined by the belief volatility parameter *ω*. Initial trustworthiness belief is estimated per participant (*μ*_0_). How beliefs about the others’ trustworthiness map on to investments is governed by the ordinal logistic link function with two additional subject-level parameters: choice uncertainty (*γ*) and the slope (*η*). The parameter estimation is done through hierarchical Bayesian inference, where we estimate all individual and group level parameters as well as drug effects on all parameters in one inferential step. **b**, Two example participants portrays the different behaviours that the model can capture. The participants have different belief volatilities for the bad trustee (*ω*_*bad*_), which determine the degree of uncertainty surrounding the trustworthiness beliefs (*σ*^(*t*)^), which in turn determines the degree of belief shifts. **c**, For each participants we randomly draw parameters from their individual posterior distribution, simulate data, and re-estimate the parameters 5 times. Relative high correlations indicate that the model parameters are well defined. **d**, The two main parameters of interest, belief volatility and choice uncertainty correlate with distinct features of behaviour.

### Genotype-dependent effects of D2/D3 receptor antagonism on belief volatility and precision-weights

For parameter estimation, we embedded the HGF derived equations in a hierarchical Bayesian model which allowed us to estimate the drug and genotype effects on all computational parameters in one inferential step (Kruschke, 2014; McElreath, 2018). Through this analysis, we found a main effect of sulpiride on volatility of beliefs (b = 0.831, 95% CrI [0.115, 1.533], P(b<0) = 0.01, d = 0.65, 95% CrI [0.088, 1.283], Fig. 3a), and an interaction effect of sulpiride with the genotype (b = -1.506, 95% CrI [-2.649, -0.411], P(b>0) = 0.004, d = -1.175, 95% CrI [-2.238, -0.306]). In fact, the effect of sulpiride on belief volatility is driven by the A1+ allele carriers (b = 1.598, 95% CrI [0.727, 2.465], d = 1.25, 95% CrI [0.533, 2.119]) and is not present in A1-group (b = 0.076, 95% CrI [-0.874, 0.985], d = 0.06, 95% CrI [-0.683, 0.783]).

**Fig. 3.**
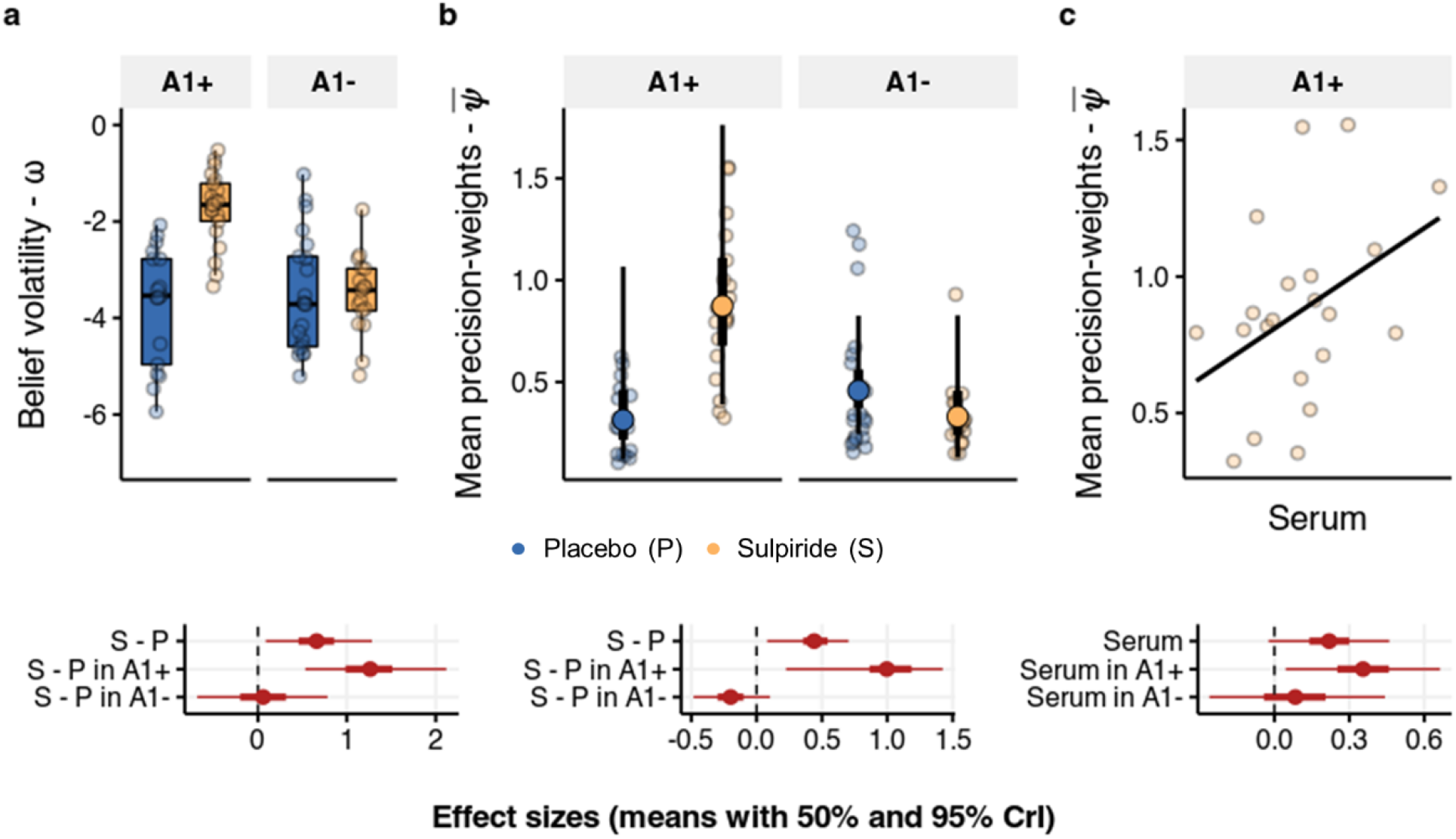
Effects of sulpiride on belief volatility and precision weights. **a**, Belief volatility boxplots over individual means of posterior distributions. Belief volatility is higher in the sulpiride group, and this effect is driven by the A1+ group (50% and 95% CrI of effect sizes below). **b** Precision-weights on PEs. Scattered points are meaned precision weights across all trials for each participant. Overlayed group level medians with 50% and 95% CrI. The effect sizes were calculated from posterior distributions of differences in means across four groups. **c**, Precision weights correlate with log transformed serum levels in the blood.

The key consequence of higher belief volatility is that it leads to lower precision of prior beliefs and therefore of predictions, which has a direct effect on the learning rates. Indeed, what we find is that participants under sulpiride have higher average precision-weights (d = 0.452, 95% CrI [0.081, 0.704], P(d<0) = 0.008, Fig. 3b), an effect again significantly more pronounced by A1+ participants (d = 1.042, 95% CrI [0.225, 1.424], P(d<0) = 0.003), and not present in the A2 homozygotes (d = -0.202, 95% CrI [-0.482, 0.103], P(d>0) = 0.089) with a significant interaction effect (d = 1.244, 95% CrI [0.335, 1.714], P(d<0) = 0.001). Importantly, in the A1+ group, this effect of sulpiride on precision-weighting correlated with the degree of serum levels in the blood (b = 0.356, 95% CrI [0.045, 0.663], P(b<0) = 0.013, Fig. 3c).

At this point we also note that there were no differences in initial beliefs (*μ*_0_) about the trustworthiness (b = -0.143, 95% CrI [-0.68, 0.389], P(b>0) = 0.301). Looking at potential asymmetries when dealing with uncertainty about positive or negative outcomes, we find that in the A1+ group the difference in *ω* between placebo in sulpiride is apparent in interactions with both trustees (b_bad_ = 2.093, 95% CrI [1.103, 3.103], P(b_bad_ <0) = 10e-3, b_good_ = 1.094, 95% CrI [0.09, 2.06], P(b_good_ <0) = 0.014), but is marginally higher for the bad trustee (b_good-bad_ = -1.009, 95% CrI [-1.962, -0.047], P(b_good-bad_ >0) = 0.021, Supplementary Fig. 4). Interestingly, this analysis also showed, that in the A1-group there is a significant interaction of sulpiride and trustee effects (b_good-bad_ = -1.696, 95% CrI [-2.758, - 0.615], P(b_good-bad_ >0) = 0.001), whereby in that genetic group the effects of sulpiride on belief volatility are marginally significant for the bad trustee (b_bad_ = 0.921, 95% CrI [-0.188, 1.995], P(b_bad_ <0) = 0.048), but not for the good trustee(b_good_ = -0.771, 95% CrI [-1.832, 0.26], P(b_good_ >0) = 0.075).

### D2/D3 receptor antagonism increases choice uncertainty

In addition to the effect on belief volatility, sulpiride also increases choice uncertainty, by decreasing the parameter *γ* (b = -1.049, 95% CrI [-1.6, -0.502], P(b<0) < 10e-3, d = -0.979, 95% CrI [-1.535, - 0.455], Fig. 4a), with smaller effects in the A1+ group (b = -0.646, 95% CrI [-1.272, -0.033], P(x>0) = 0.02, d = -0.608, 95% CrI [-1.206, -0.031]) and more prominent effects in the A2 group (b = -1.44, 95% CrI [-2.261, -0.639], P(b<0) = 10e-3, d = -1.351, 95% CrI [-2.133, -0.601]). Since lower values of *γ* correlated with higher proportion of mistake trials we examined how sulpiride affected the proportion of mistake trials and found that it on average increased the number of mistake trials (b_logodds_ = 1.172, 95% CrI [0.443, 1.992], P(b_logodds_ <0) < 10e-3, Fig. 4b), an effect driven by the A1-group (b_logodds_ = 1.876, 95% CrI [0.781, 3.032], P(b_logodds_ <0) < 10e-3) and was not present in the A1+ group (b_logodds_ = 0.468, 95% CrI [-0.535, 1.537], P(b_logodds_ <0) = 0.184), with a marginally significant interaction (b_logodds_ = 0.885, 95% CrI [-0.041, 1.857], P(b_logodds_ <0) = 0.03). The effect of sulpiride on the proportion of mistake trials in the A1-group was proportional to the blood serum levels (b_logodds_ = 0.607, 95% CrI [0.089, 1.142], P(b_logodds_ <0) = 0.011) with no correlation of the A1+ group (b_logodds_ = -0.328, 95% CrI [-0.818, 0.143], P(b_logodds_ >0) = 0.085). This parameter also determines the skew in the distribution of investments, whereby higher values make extreme investments more likely (Fig. 4c, d). At this stage we also note that from a perspective of an expected utility maximising agent, extreme investments are most optimal (Supplementary Note 2). Individuals with higher *γ* therefore behave more as rational agents and take the uncertainty of the outcome less into consideration when choosing investments. Sulpiride also increased the *η* parameter (b = 1.459, 95% CrI [0.532, 2.42], P(b<0) < 10e-3, d = 0.941, 95% CrI [0.331, 1.58]), further advocating for the assertion that sulpiride increased the degree to which beliefs about trustworthiness influenced participants’ investments.

**Fig. 4.**
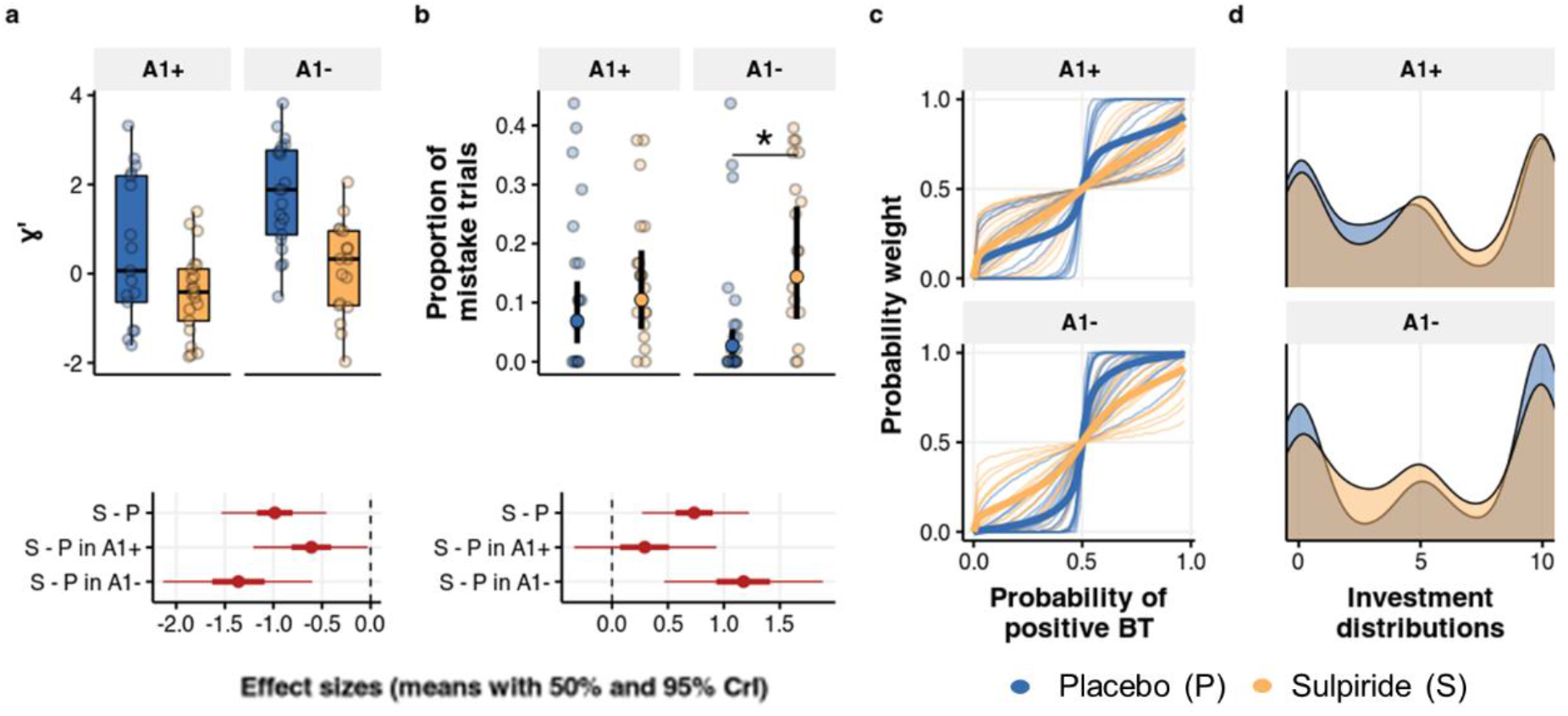
Effect of sulpiride on choice uncertainty. **a**, Effects of sulpiride on choice uncertainty, estimated in log – space (hence the prime). Dots are participant-level posterior means. The 95% and 50% CrI of effect sizes show a main effect of sulpiride, driven pa the A1-group. **b**, Proportion of mistake trials is higher following sulpiride administration in the A1-group, but not in the A1+ group. Means and 95% quantiles of posterior distributions across the four groups are plotted based on a logistic regression model. Corresponding effect sizes below. **c**-**d**, The choice uncertainty parameter determines the probability weight (**c**) and therefore the investment distribution (**d**). Higher values for the placebo group (the A1 group in particular) indicate more extreme investment choices and higher belief inflexibility.

In sum, the overall results from the computational modelling suggest that sulpiride treatment led to higher choice uncertainty, which was related to increased mistakes in the in the A1-group specifically. Sulpiride also increased belief volatility and precision-weights on PEs, an effect that was driven by the A1+ group, and in the A1-group was only marginally present for the bad trustee, but not present overall. An important final step was to exclude the possibility that this increase in updating was due to increased sensitivity to social feedback in general, or due to decreased desire to maximise outcomes. For this, we turned to data from single-round social interaction games that measure learning-independent positive and negative reciprocity.

### No effect of D2/D3 receptor antagonism on single-round reciprocal behaviour

In the single-round interaction games the participants played a slightly modified versions of the trust game. In the positive reciprocity game, they played the trustee and could reward the investor for their decision (Fig. 5a). In the negative reciprocity game, they played as investor and could punish the trustee (Fig. 5b). We found no differences between sulpiride and placebo, neither in the amount of reward (Back-transfer) in the positive reciprocity game (b = -0.023, 95% CrI [-6.605, 6.263], d = 0.000, 95% CrI [-0.032, 0.03], Fig. 5c) nor in punishing behaviour in the negative reciprocity game (b = 1.552, 95% CrI [-0.903, 3.98], P(x<0) = 0.106, d = 0.2, 95% CrI [-0.114, 0.513], Fig. 5d). This implies that the drug effect on reciprocal behaviour in the Repeated Trust Game was not due to higher sensitivity to social-feedback, or to less rational behaviour.

**Fig. 5.**
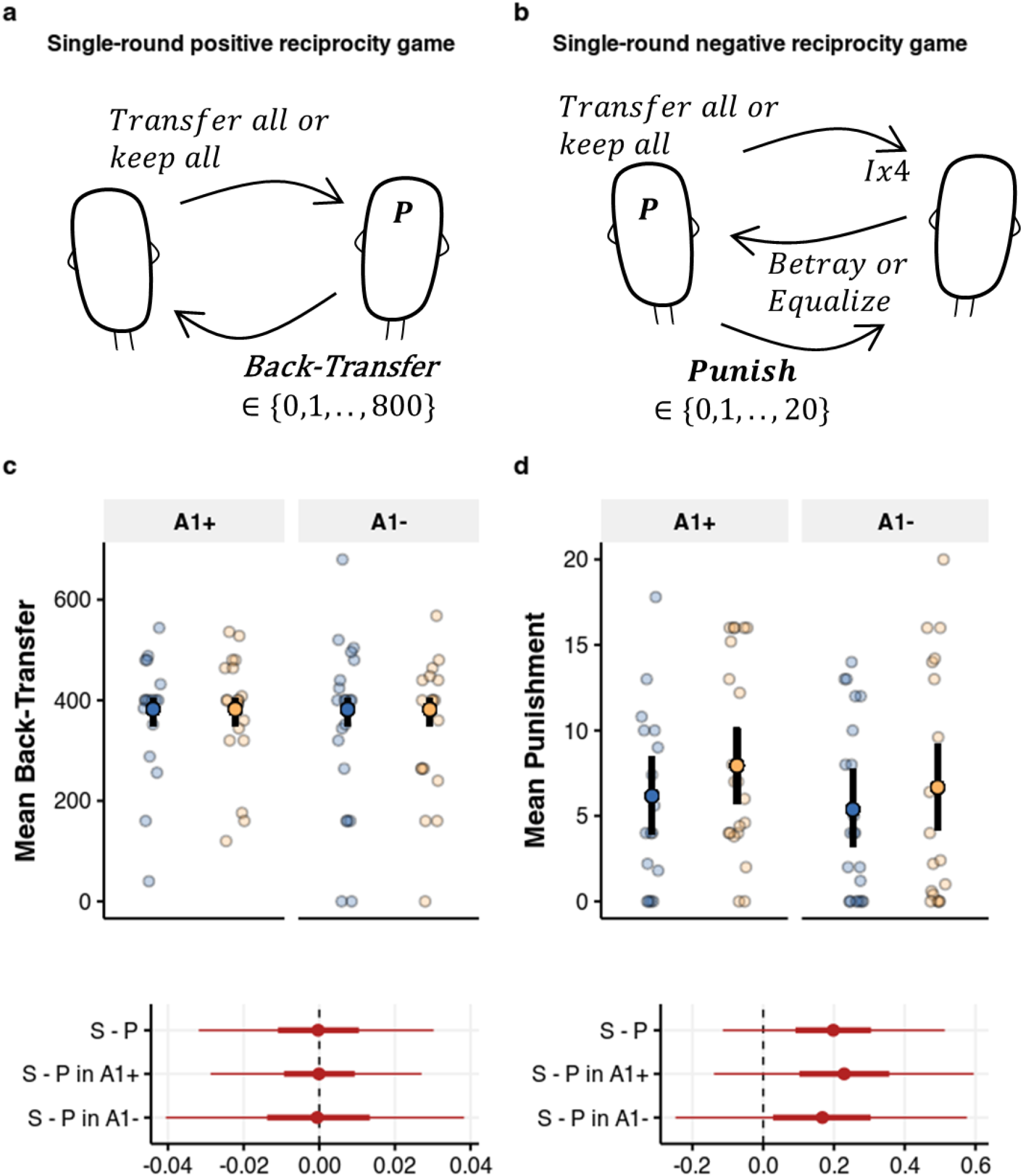
Single-round reciprocity games. **a**, In the positive reciprocity task, participants played as trustees were paired with 7 other players. The investor in this version of the game received 800 points and could either keep everything or give everything to the trustee, who could then decide how to split the points. The investors were pre-programmed so that 5 out of 7 transferred everything to the trustee. **b**, In the single-round negative reciprocity game, the participants played the investor. In the beginning of the round both players were given 10 points. The investor could then decide to transfer everything or nothing. The transferred investment got multiplied by a factor of four and the trustee could them decide to either equalize or betray. Crucially, after the choice of the trustee, both players received another 20 points and the investor could use his points to punish the trustee, with a factor of three. **c, d**, Mean and 95% CrI of Back-transfer (**c**) and punishment (**d**) across the four groups, plotted over raw means per participant, with no differences across groups, as evident from the means and 95% CrI of effect sizes shown below.

## Discussion

Inferring attitudes of others is fundamental to our social functioning, but the neurocomputational mechanisms of the updating of beliefs about others are not well understood. We show that blocking D2/D3 dopamine receptors has a profound effect on how healthy participants process uncertainty in a social context. When playing as investors in the Repeated Trust Game, participants given a high dose of sulpiride changed their investment more from one trial to the next. Using a hierarchical Gaussian filter to explicitly model the evolution of participants’ beliefs about the trustworthiness of the trustees, we show that sulpiride increased belief volatility. This implies that for the participants under sulpiride, the beliefs about the trustworthiness of others were held with less precision (i.e., with higher uncertainty), which in turn caused increased precision weights on PEs. This effect was more pronounced in participants with at least one minor A1 allele of the Taq1a polymorphism, associated with higher endogenous dopamine levels. The increase in precision weights on PEs in that genetic subgroup scaled with the sulpiride serum levels in the blood. As a consequence, sulpiride led to higher reciprocal behaviour (increased investment after negative back-transfer and decreased investment after positive back-transfer), but in the repeated trust game and not when interacting with individuals in single-round interactions. Moreover, sulpiride decreased the value of the parameter of the model that codes for deterministic action selection policies (*γ*), implying higher uncertainty about investment selection. The effect was present in both genotype groups.

On the neurophysiological level, it has been proposed that precision is encoded through the post-synaptic gain (i.e. amplification or stifling of presynaptic neuronal input) of neurons that propagate PE signals (Friston et al., 2012). Our results are in-line with the idea that dopamine binding to D1 receptors of the medium spiny neurons in the striatum increases the gain on PE signals, while binding to D2 receptors decreases gain through disinhibition of the so called indirect pathway (Frank, 2005; Yao et al., 2008). Within this framework increased precision-weights following D2 antagonism can be explained by more dopamine being available to bind to D1-like receptors, a claim that is further substantiated by the observation that the effect of sulpiride was stronger in participants with genetically conferring higher presynaptic dopamine availability and lower D2 receptor density. These findings extend previous studies highlighting the role of dopamine receptors in coding precision or uncertainty in various contexts, such as perceptual and risk-based decision making (Guggenmos, Wilbertz, Hebart, & Sterzer, 2016; Schwartenbeck et al., 2015). In particular, previous work has shown that sulpiride decreased the perceived precision of temporal expectations (Tomassini, Ruge, Galea, Penny, & Bestmann, 2016). In a task where participants were explicitly told about the variance of outcomes, they adapted their behaviour accordingly, which led to more optimal choice performance (Diederen, Spencer, Vestergaard, Fletcher, & Schultz, 2016). This behavioural pattern was accompanied by adaptive PE signals in the midbrain and the ventral striatum. Under 600 mg of sulpiride, both the PE scaling as well as the adaptive PE coding in the midbrain and partially in the striatum were reduced (Diederen et al., 2017). This suggests that D2 receptors likely play a general role in uncertainty coding across various task modalities and contexts.

Our findings that blocking D2/D3 receptors increases learning rates may seem to be at odds with previous work showing that D2/D3 antagonists reduced performance in learning tasks and attenuated prediction error signals in the striatum (Jocham et al., 2014; Pessiglione et al., 2006) as well as with studies showing no effect of D2/D3 antagonism on learning rates (Jocham et al., 2011, 2014), even when using similar computational frameworks (Iglesias et al., 2021; Marshall et al., 2016). It is thus important to consider that the A1 is considered a minor allele of the Taq1a polymorphism, meaning that in most other studies participants were likely predominantly A2 homozygotes. We observed a more general effect of D2/D3 receptor antagonism on choice uncertainty that was more prominent in A2 homozygotes and was related to a higher number of mistake trials in that subgroup of participants, although the number of mistakes was not high enough to reduce investment on average. Furthermore, participants could invest on an ordinal 11-point scale, which allowed us to capture smaller belief shifts that might either be missed in learning tasks with categorical choice options or be attributed to a different choice selection policy. For example, the participants in our study also performed a standard probabilistic two-bandit task afterwards, where participants in the A1+ group under sulpiride compared to placebo continued to switch between choice options, which was explained by increased choice stochasticity, parametrized through the soft-max decision temperature (Eisenegger et al., 2014).

The increase in choice uncertainty or stochasticity under sulpiride that we observed, could have been due to participants being less motivated to maximise outcome, and therefore less likely to behave as a rational “homo economicus” (Camerer, 2003). D2 receptors do play a role in motivation. For example, optogenetic excitation and inhibition of D2 receptors in the ventral striatum of rats is reported to respectively increase and decrease motivation (Soares-Cunha et al., 2016). Were that the case in our study, one would expect a different behavioural pattern in the single shot trust games, where rational agents would not punish betrayals nor reward trusting behaviour (Fehr, Fischbacher, & Gächter, 2002), which we did not. Alternatively, it is possible that the increased action variability resulted from noise in belief updating and not in choice selection per se (Findling, Skvortsova, Dromnelle, Palminteri, & Wyart, 2019). What speaks against this interpretation is that the overall performance in the task was not reduced following sulpiride administration for neither of the genetic subgroups, suggesting that the investment selection under sulpiride was not random, but only less precise.

Indeed, variability in investment selection following sulpiride administration is well in line with what we know about the role of dopamine receptors in action selection. In particular, whereby the D1 receptor expressing cells on the direct pathway facilitate action execution, the D2 receptor expressing cells on the indirect pathway suppress action selection (Frank, 2005). Stimulation of D2 receptors through endogenous dopamine leads to inhibition of the indirect pathway and increases the probability of repeating the same action. Accordingly, blockade of postsynaptic D2 receptors increases the probability of performing competing actions and therefore promotes randomness in action selection (Sridharan, Prashanth, & Chakravarthy, 2006). For example in macaques, microinfusion of D2 (but not D1) receptor antagonists into the dorsal striatum led to increased choice stochasticity (Lee, Seo, Dal Monte, & Averbeck, 2015) and a similar pattern was observed in D2 receptor knockout mice (Kwak, Huh, Seo, Lee, & Han, 2014). In humans, a recent positron emission tomographic imaging study showed that D2 receptor availability in the striatum correlated with deterministic decision-making strategies represented either through decision temperature within reinforcement learning as well as with policy precision within active inference (Adams et al., 2020).

The key idea of active inference models is to extend the Bayesian generative models of beliefs about the states of the world, to include beliefs about preferred states, therefore casting both action and perception as an inference problem (Friston et al., 2015, 2013). An active inference agent thus prefers actions that minimize the statistical distance (relative entropy) between the distributions of desired and predicted future states. The expected precision of a policy, in the context of our task, controls the confidence with which the participants selected a certain action, which can explain the more variable investment we observed in the sulpiride group. Within this framework, we can interpret the effects of sulpiride in our study as reflecting a more general role of D2 receptors in coding precision of both beliefs and action policies, thus extending previous theoretical and experimental work on the involvement of dopamine in modulating precision in predictive coding schemes (Friston et al., 2012; Nour et al., 2018; Schwartenbeck et al., 2015).

Our findings are particularly relevant for understanding the effects of antipsychotic medication in patients with psychosis, a disorder characterized by rigid beliefs of persecution, underlined by a profound lack of trust in others (Freeman, 2016; Fuchs, 2015). Previous studies with repeated trust games showed that patients with psychosis have lower initial trust and find it hard to change their beliefs (Fett et al., 2012; Gromann et al., 2013). Neurocomputational accounts of delusions suggest that patients are impaired in forming higher level models and making accurate predictions (Adams et al., 2013; Sterzer et al., 2018), leading to a world that is constantly surprising and full of salient events that need explanation (Kapur, 2003). As a consequence, patients rely more on internal sources of information (Schmack, Schnack, Priller, & Sterzer, 2015; Teufel et al., 2015) and negative self-schemas (Bentall, Kaney, & Dewey, 1991; Fuchs, 2015; Rossi-Goldthorpe, Leong, Leptourgos, & Corlett, 2021), leading them to “jump to conclusions” (Garety, Hemsley, & Wessely, 1991; Van Der Leer & McKay, 2014), and form rigid convictions held with high confidence (Moritz et al., 2014; Rossi-Goldthorpe et al., 2021; Woodward, Moritz, Menon, & Klinge, 2008). In our data, acute administration of an antipsychotic increased belief volatility and reduced deterministic choice strategies. Our findings thus support the view that antipsychotics help to reduce the impact of distressing beliefs and therefore offer a therapeutic window within which adjunct therapies might be more efficient in helping the patients to resolve their issues (Bentall, Kinderman, & Kaney, 1994; Hole, Rush, & Beck, 1979; Kapur, 2003). It is however up to future studies to directly test this proposal in patients.

In conclusion, we show that blocking D2 dopamine receptors increases the flexibility of beliefs when learning about others. This finding importantly contributes to our understanding of how the brain infers the attitudes of other people. By mapping out the connection between alterations in the dopaminergic system with specific computational substrates this study not only contributes to the advancement of our knowledge of how the brain performs inference, but also to our understanding of when it fails appropriately to do so.

## Acknowledgements

This paper is dedicated to the memory of Christoph Eisenegger. This research work was funded by a Core Award from the Medical Research Council to the Behavioural and Clinical Neuroscience Institute (MRC Ref G1000183; WT Ref 093875/Z/10/Z) and the Vienna Science and Technology Fund (WWTF) with a grant (VRG13-007) awarded to Christoph Eisenegger and Claus Lamm. Christoph Eisenegger was also supported by the Swiss National Science Foundation (PA00P1_134135). We are grateful for the participation of all NIHR Cambridge BioResource (CBR) volunteers and thank the Cambridge BioResource staff for their help with volunteer recruitment. We also thank members of the Cambridge BioResource SAB and Management Committee for their support given to our study and the National Institute for Health Research Cambridge Biomedical Research Centre for funding.

## Conflict of interest

LC has received royalties from Cambridge Cognition Ltd. relating to neurocognitive testing. UM discloses consultancy for Janssen-Cilag, Lilly, Heptares and Shire, and educational funding from AstraZeneca, Bristol-Myers Squibb, Janssen-Cilag, Lilly, Lundbeck and Pharmacia-Upjohn. TWR discloses consultancy with Cambridge Cognition Ltd and a research grant with Shionogi Inc. The remaining authors declare no conflict of interest.

## Methods

The data and analysis scripts are available online (https://github.com/nacemikus/belief-volatility-da-trustgame.git).

### Participants

Data were collected from 78 male participants, aged between 19 and 44 years (mean= 32.1), recruited from a large panel of volunteers, that were genotyped and screened for mental and physical health (Cambridge BioResource). Only participants with no history of neurological or psychiatric disorder were included in the study. Participants were stratified based on the Taq1A genotype into two groups: participants carrying at least one A1 allele (A1+), and A2 allele homozygotes (A1-). The study was performed in accordance with the Declaration of Helsinki and approved by the National Research Ethics Committee of Hertfordshire (11/EE/0111).

### Procedure

After arrival participants underwent another psychiatric screening and an alcohol test to exclude alcohol consummation on the study day. After an assessment of general intelligence (National Adult Reading Test) participants signed an informed consent before they were administered a single oral dose of either 800 mg of sulpiride or placebo in a randomized, double-blind fashion. We used the parallel group design, because complex behavioural tasks (like the trust game) have practice (repetition) effects that can confound the results of within group pharmacological experiments. Before behavioural testing participants waited for three hours in a quiet room, where they were allowed to read a newspaper. To monitor the effects of the pharmacological manipulation, blood pressure and heart rate and mood and drug effects were assessed prior to drug administration and after the waiting period. Similarly, blood samples to determine the serum levels were taken at both time points. The social interaction tasks presented here included a repeated trust game, and positive and negative reciprocity tasks and were part of a test battery that included a working memory task and an instrumental learning task, both published elsewhere (Eisenegger et al., 2014; Naef et al., 2017). Two participants were excluded from the analysis: one felt uncomfortable in the room, and one did not sufficiently understand the instructions of the social interaction tasks. This led to the following group distributions: 17 A1 allele carriers received placebo, and 21 received sulpiride, and 21 A2 homozygotes received placebo, and 17 received sulpiride. Participants were matched across the four groups for age, body mass index, general and verbal intelligence (Table 1, all p>0.30). Participants received a monetary compensation of £50 plus the extra money earned in the behavioural tasks.

**Table 1.** Demographic information.

| Genotype | Treatment | N | IQ | sd | Verbal IQ | sd | BMI | sd |
| --- | --- | --- | --- | --- | --- | --- | --- | --- |
| A1+ | Placebo | 17 | 120.2 | 5.333 | 120.335 | 5.913 | 26.098 | 3.125 |
| A1+ | Sulpiride | 21 | 120.21 | 7.334 | 120.355 | 8.133 | 26.595 | 5.74 |
| A1- | Placebo | 21 | 120.05 | 6.089 | 120.17 | 6.76 | 24.604 | 4.415 |
| A1- | Sulpiride | 17 | 117.659 | 7.494 | 117.518 | 8.318 | 25.064 | 5.393 |

sd, standard deviation

### Sulpiride Serum Concentration Measurements

The level of serum sulpiride was determined by high-performance liquid chromatography. This method utilizes fluorescence endpoint detection with prior solvent extraction. The excitation and emission wavelengths were 300 and 360 nm, respectively. Both intra- and inter-assay coefficients of variation (CVs) were 10% and the limit of detection was 5–10 ng/ml.

### Prolactin level assessment

The prolactin level was measured using a commercial immunoradiometric assay (MP Biomedicals, Santa Ana, CA, USA), 3h after capsule ingestion. Prolactin levels were expected to increase with blocking postsynaptic D2 receptors (Wiesel, Alfredsson, Ehrnebo, & Sedvall, 1982). The intra- and inter-assay coefficients of variation were 4.2% and 8.2%, respectively, and the limit of detection was 0.5 ng ml−1 (Eisenegger et al., 2014). We found that sulpiride administration significantly increased blood plasma prolactin levels (Δ = 33.1 mg/ml, p < 10e-3), and this increase was significantly higher (p < 10e-3) than the changes in the placebo group (Δ = -0.91 mg/ml, Mann Whitney test for differences p < 10e-3). Data for three volunteers were excluded due to blood contamination.

### Side-effects and mood assessments

Side effects were assessed with a neurovegetative list (Rush, Stoops, Hays, Glaser, & Hays, 2003), 3 h after drug intake. Mood was assessed with a visual analogue scale at baseline and 3 h after drug intake. Items in the visual analogue scales (VAS) were alert/drowsy, calm/excited, strong/feeble, muzzy/clear-headed, well coordinated/clumsy, lethargic/energetic, contented–discontented, troubled–tranquil, mentally slow/quick-witted, tense/relaxed, attentive/dreamy, incompetent/proficient, happy/sad, antagonistic/amicable, interested/bored and withdrawn/gregarious. The factors “alertness”, “contentedness”, and “calmness” were calculated from these items (Eisenegger et al., 2010). Data from one volunteer were excluded due to technical issues. We found no apparent drug effects on mood, heart rate, blood pressure or self-reported side-effects (for details see Supplementary Material).

### Repeated trust game

In the trust game (Berg et al., 1995) an investor (Player A) decides on how much money they want to transfer to the other player, called the trustee (Player B). The trustee receives the investment that is however tripled by the experimenter and decides on how to split the acquired sum. We used a multi-round version of the task (King-Casas, 2005), where the interchange between the investor and the trustee repeated across 25 trials. In the beginning of each trial both players were endowed with 10 points, to avoid investments motivated by inequality aversion. Each point converted to two pence at the end of the experiment. The participants could invest points on a scale from 0 to 10 and the trustee could respond in a binary fashion, by either equalizing the payoff, or defecting by keeping all the points in the trial for themselves. Participants played as investors against two pre-programmed trustees: one defected in 7 out of 25 trials (the good agent) and the other defected in 18 out of 25 trials (the bad agent). To increase ecological validity, the participants were led to believe that they play against two actual people that have already given their answers in advance several weeks before the testing, and that their decision will impact the payoff of these participants.

### Positive and negative reciprocity games

In the positive reciprocity game, the two players need to distribute 800 points. First, player A is offered a distribution whereby they get 800 points and player B gets 0 points. They can decide to either keep all the points or delegate the decision on how to divide the points to player B. If the decision was to delegate, to player B can decide on any point distribution between the two players. Participants in our study played as player B sequentially against 7 different people playing as player A. The negative reciprocity game is like a trust game in which defecting behaviour of the trustee can be punished by the investor. Both players are first endowed with 10 points. Player A then decides to either transfer his endowment (all the 10 points) all transfer nothing. The transfer of player A is quadrupled by the experimenter. Player B can then decide to either keep everything to themselves or to equalize the payoff. Following the decision of player B, both players get endowed with another 20 points and player A can spend each of these 20 points to penalize player B’s outcome, whereby each penalty point of player A spent this way deducts three times as many points from player B’s outcome. Participants in our study played as player A against 7 different people playing as player B. The actions of people playing player B were pre-programmed so that 5 out of 7 defected.

In both games the participants were told that the players have given their answers already days before the testing. Each point converts to 0.2 pence for the positive reciprocity game and 4 pence for the negative reciprocity game.

### Behavioural analysis

Behavioural analysis was done with Bayesian multilevel (generalized) linear regression (McElreath, 2018), fitted with the brms package in R (Bürkner, 2017). All models were run with 4 chains, 3000 iterations each with 800 warmup. The quality of chain convergence was inspected visually based on trace plots of main fixed effects, and a threshold on Gelman-Rubin 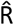 Statistic for each parameter was set to 1.01 (Gelman & Rubin, 1992). Throughout the behavioural analysis we z-scored the dependent variables, coded the Treatment variable as 0 (placebo) and 1 (sulpiride), Back-transfer as 1 (equalize) and -1 (betray), Genotype as 0 (A1-) and 1 (A1+) and centred the trustee variable (0.5 for good, and - 0.5 for bad trustee). All random effects were modelled as a multivariate normal distribution, thereby evaluating the correlation between the effects as well as pooling information across the effects. Priors used are depicted in Table 2. The effect sizes where calculated by dividing the regression coefficients with the square root of summed variances of the residuals and of all random effects (Nalborczyk, Batailler, Loevenbruck, Vilain, & Bürkner, 2019). All models were redone also in the *lme4* package (Bates, Mächler, Bolker, & Walker, 2014) or *nlme* package (Pinheiro, Bates, DebRoy, Sarkar, & Team, 2007), and the results of those models are reported in the supplementary material.

**Table 2:**
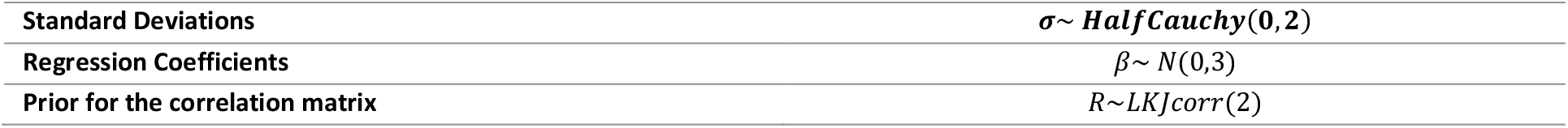
Prior distributions for the analysis of investment changes.

#### Analysing investment behaviour

All model summary tables are in the supplementary material. The effect of sulpiride on absolute change from one trial to the next was evaluated with a model predicting effects on absolute change of sulpiride and trials with random intercepts for each participant. We also rerun the model including a participant-level slope for the trial and found that it does not affect inference about the main effect, but does increase the uncertainty around the interaction term. Next, the Genotype and Trustee as group level predictors were included as well as a random slope for the Trustee for each participant. Since the dependent variable is bounded at 0, the same analysis was done again with the dependent variable shifted by 1 and log transformed. This did not affect the conclusion of the model.

To analyse relative changes in investment the z-scored relative change from one trial to the next was predicted from the variable for Back-transfer (coded as -1 and 1), Treatment, Genotype, Trustee as well as their interactions, with a participant-level random intercept and slope for the Trustee.

The reciprocal and mistake trials were analysed with a multilevel logistic regression model including predictors Treatment, Genotype, Trustee, and their interaction, again with a random intercept and slope for the effects of the Trustee for each participant.

To analyse average investments, we used a multilevel ordinal-logistic regression model, with Treatment, Genotype, Trustee, Trial and their interaction, with a random intercept and slope for the effects of the Trustee for each participant.

Models analysing the single-round reciprocity tasks predicted Punishment (negative reciprocity) and Back transfer (positive reciprocity) from Treatment, Genotype and their interaction, including a random intercept per participant.

### Computational modelling

We first defined a generative model of the evolution of beliefs about the other players’ trustworthiness as a Gaussian random walk. The belief volatility parameter *ω* describes the degree to which these beliefs can change from one trial to the next. We then used the Hierarchical Gaussian Filter (HGF) to invert this model (Mathys et al., 2011).

#### Generative Model

The generative model describes the evolution of beliefs about the other person’s trustworthiness as a Gaussian random walk with a step size of exp(*ω*). In particular, at trial *t* the belief on the other player’s trustworthiness is defined as

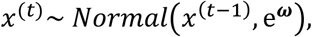

where *ω* is a participant level parameter. The mapping from the trustworthiness beliefs to the probability of a positive Back transfer (*BT*) occurs through a sigmoid transform *s*(.). So, at trial *t* we define:

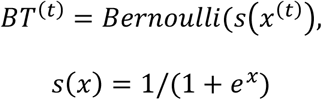

#### Model inversion and update equations

To define the inferred participant level belief trajectories the generative model is inverted using Hierarchical Gaussian Filtering (Mathys et al., 2011, 2014). The HGF approximates full Bayesian inference using variational Bayes to derive at trial level update equations that resemble those of a Kalman filter (Gershman, 2015; Kakade & Dayan, 2002). In particular, the weights (learning rates) on the PEs are determined by the precision of prior beliefs as well as the uncertainty about the outcome. The HGF provides inferred posterior distributions of participants’ belief trajectories as Gaussians through the mean *μ*^(*t*)^ and variance 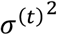 or its inverse, the precision *π*^(*t*)^ in the update equations for both time series:

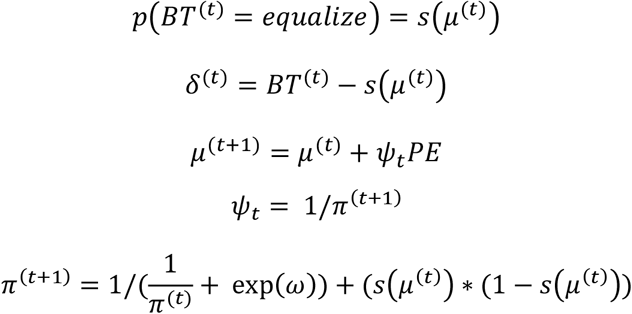

This is a so-called recognition or perceptual model and describes our beliefs about the belief of the participants. To map these beliefs on to the behaviour of the participant a response model is defined through a likelihood function. Because investments occur on an ordinal scale we used the ordered logistic link function (Bürkner & Vuorre, 2019):

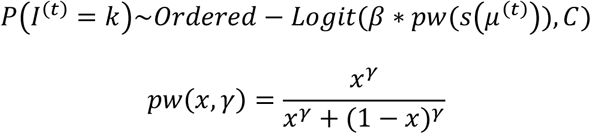

Where *C* is a vector of intercepts and *pw*(.) is a probability weighting function on the unit interval. The ordered-logit estimates 10 intercepts, that determine the mapping from the linear term to the ordinal investments. In an ideal case, we would estimate all 10 intercepts for each subject, which was not feasible with our data. We therefore estimate the 10 intercepts for all subjects, add one subject-level intercept in the linear term and assist the model in accounting for the various investment distributions with the probability weighting (*pw*). We compared this model to an HGF model without the probability weighting function and a simple Rescorla-Wagner model (Rescorla & Wagner, 1972) with separate learning rates for positive and negative feedback.

#### Parameter Estimation

The model parameters were estimated in one hierarchical Bayesian model. This approach reduces overfitting (McElreath, 2018), pools information across different levels (drug groups, and participants) and allows us to estimate both participant and group level parameters in one inferential step. Meaning, we estimate the effects of our drug manipulation on all relevant computational parameters in one model, while at the same time, leading to more stable parameter estimates (Ahn, Haines, & Zhang, 2017). Models were implemented in Stan (Carpenter et al., 2017) using R as the interface. Each candidate model with four independent chains and 3200 iterations (800 warm-up). Convergence of sampling chains was estimated through the Gelman-Rubin 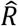 statistic (Gelman & Rubin, 1992), whereby we considered 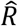 values smaller than or equal to 1.01 as acceptable.

The intercepts from the response model, *c*_*k*_, *k=* 1, …, 10, were estimated on the group level. This determined a general mapping from the probability to Investment. The participant level parameters (*ω*, Δ*ω, γ, η* and *μ*_0_) were modelled as a multivariate Gaussian distribution:

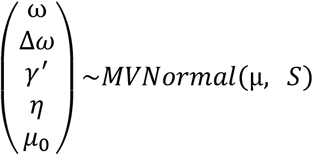

Where *S* is the covariance matrix and *μ* is the vector of means. The Δ*ω* parameter denotes the modelled difference in *ω* between the good and the bad Trustee. The matrix *S* was factored into a diagonal matrix with standard deviations and the correlation matrix *R* (Bürkner, 2017; McElreath, 2018). The prime denotes the parameters in estimation space, whereby *γ* was estimated in log space, due to it being lower bound by 0. The vector *μ* included all group level regression coefficients for the drug, genotype, and their interaction. Meaning, that participant level *ω* parameters were drawn from a distribution with the following mean:

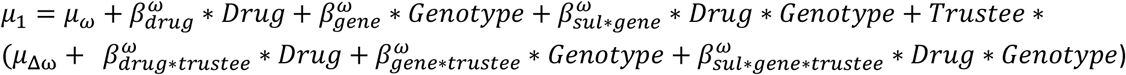

And the means of the rest of the parameters were defined as:

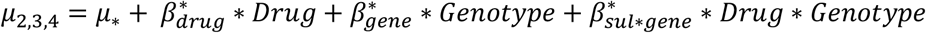

Using the following priors

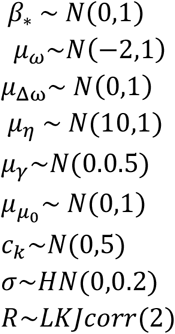

where 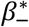 are coefficients, drawn from *N*(0,1). The priors for group-level means for non-transformed parameters were weakly informative, for *γ*, estimated in log-space, the prior was *N*(0,0.5), the prior for group-level standard deviations were more regularizing, with *σ* ∼*HalfNormal*(0,0.2), and the prior for the correlation matrix was *R*∼*LKJcorr*(2). The prior for group level *η* was set to something above 0, because chains that sampled from areas too close to 0 usually got stuck in that area.

### Model Validation and Comparison

For parameter recovery 5 parameter sets were drawn from each participant’s mean and standard deviation and used to simulate data. Simulated data were then estimated with the same model and the re-estimated parameters were correlated with the simulated ones. Further, posterior distributions of parameters were used to simulate data and check whether the crucial aspects of behaviour are captured by the model. A trial based Leave-One-Out Information Criterion (LOOIC) was used to compare the three models (Vehtari, Gelman, & Gabry, 2017) using the loo package in R. The LOOIC approximates out-of-sample predictive accuracy of each trial, with lower LOOIC scores indicating better prediction accuracy out of sample.

## Notes

https://github.com/nacemikus/belief-volatility-da-trustgame.git

