## Supplementary Material for "Blocking D2/D3 dopamine receptors increases volatility of beliefs when we learn to trust others"

### Supplementary Tables

| Abs_change~Treatment*Trial + (1\|ID) | Estimate | Est.Error | Q2.5 | Q97.5 |
| --- | --- | --- | --- | --- |
| **Intercept** | -0.15 | 0.076 | -0.295 | -0.003 |
| **TreatmentSulpiride** | 0.309 | 0.106 | 0.108 | 0.518 |
| **Trial** | -0.015 | 0.003 | -0.021 | -0.009 |
| **TreatmentSulpride:Trial** | 0.007 | 0.004 | -0.001 | 0.015 |
| Abs_change~Treatment*Trial + (Trial\|ID) |  |  |  |  |
| **Intercept** | -0.114 | 0.069 | -0.246 | 0.025 |
| **Treatmentsulpiride** | 0.234 | 0.097 | 0.043 | 0.427 |
| **Trial_c** | -0.012 | 0.005 | -0.022 | -0.001 |
| **Treatmentsulpiride:Trial_c** | 0.007 | 0.007 | -0.008 | 0.022 |

Supplementary Table 1: Results the model Bayesian mixed model predicting absolute changes in investment from one trial to the next including drug treatment and trial as predictors. Similar results are achieved when the slope for the trial is allowed to vary per participants. Treatment coded as 1 sulpiride and 0 placebo, trial used as a continuous variable and centred. Refer to the code about the effect in native space and calculation of effect sizes.

| Abs_change ~Treatment*Trial + (1\|ID) | Value | Std.Error | DF | t-value | p-value |
| --- | --- | --- | --- | --- | --- |
| **(Intercept)** | -0.113 | 0.066 | 3570 | -1.715 | 0.086 |
| **TreatmentSulpiride** | **0.235** | **0.093** | **74** | **2.515** | **0.014** |
| **Trial** | -0.012 | 0.003 | 3570 | -3.87 | <10e3 |
| **TreatmentSulpiride:Trial** | 0.007 | 0.004 | 3570 | 1.626 | 0.104 |

Supplementary Table 2: The output of the non-Bayesian absolute change model.

| log(1+Abs_change) ~ Treatment*Trial + (1\|ID) | Value | Std.Error | DF | t-value | p-value |
| --- | --- | --- | --- | --- | --- |
| **(Intercept)** | -0.149 | 0.073 | 3570 | -2.045 | 0.041 |
| **Treatmentsulpiride** | **0.309** | **0.103** | **74** | **3.004** | **0.004** |
| **Trial** | -0.015 | 0.003 | 3570 | -5.084 | <10e3 |
| **Treatmentsulpiride:Trial** | 0.007 | 0.004 | 3570 | 1.715 | 0.086 |

Supplementary Table 3: The non-Bayesian absolute change model reran again with a transformed outcome variable. Because the absolute change variable is positive, we use a log transform of a translation.

| Abs_change ~ Treatment*Genotype *Trustee *Trial + (Trustee \|ID) | Estimate | Est.Error | Q2.5 | Q97.5 |
| --- | --- | --- | --- | --- |
| **Intercept** | -0.109 | 0.069 | -0.25 | 0.024 |
| **Treatmentsulpride** | **0.234** | **0.099** | **0.044** | **0.431** |
| **Trial** | **-0.013** | **0.003** | **-0.018** | **-0.007** |
| **Trustee** | 0.003 | 0.07 | -0.133 | 0.143 |
| **Genotype** | 0.079 | 0.138 | -0.195 | 0.348 |
| **Treatmentsulpride:Trial** | 0.007 | 0.004 | -0.001 | 0.016 |
| **Treatmentsulpride:Trustee** | -0.145 | 0.1 | -0.339 | 0.046 |
| **Trial:Trustee** | **0.015** | **0.006** | **0.003** | **0.027** |
| **Treatmentsulpride:Genotype** | **-0.111** | **0.198** | **-0.494** | **0.278** |
| **Trial:Genotype** | -0.011 | 0.006 | -0.023 | 0 |
| **Trustee:Genotype** | -0.002 | 0.137 | -0.272 | 0.274 |
| **Treatmentsulpride:Trial:Trustee** | -0.007 | 0.009 | -0.024 | 0.01 |
| **Treatmentsulpride:Trial:Genotype** | **0.021** | **0.009** | **0.004** | **0.038** |
| **Treatmentsulpride:Trustee:Genotype** | -0.051 | 0.194 | -0.433 | 0.332 |
| **Trial:Trustee:Genotype** | -0.016 | 0.012 | -0.04 | 0.008 |
| **Treatmentsulpride:Trial:Trustee:Genotype** | 0.042 | 0.017 | 0.009 | 0.076 |

Supplementary Table 4: Bayesian mixed model predicting absolute changes in investment from one trial to the next including drug treatment, trial, genotype and trustee as predictors. Treatment coded as 1 sulpiride and 0 placebo, trial used as a continuous variable. Variables Trial, Genotype (0.5 A1+, -0.5 A1-) and Trustee (0.5 good, -0.5 bad) were centred.

| Change ~Treatment*Genotype*Backtransfer*Trustee+ (Trustee \|ID) | Value | Std.Error | DF | t-value | p-value |
| --- | --- | --- | --- | --- | --- |
| **(Intercept)** | 0.017 | 0.038 | 3560 | 0.436 | 0.663 |
| **Treatmentsulpiride** | -0.002 | 0.051 | 72 | -0.034 | 0.973 |
| **Backtransfer** | 0.204 | 0.038 | 3560 | 5.337 | <10e3 |
| **GenotypeA1-** | -0.033 | 0.051 | 72 | -0.65 | 0.518 |
| **Trustee** | -0.105 | 0.076 | 3560 | -1.375 | 0.169 |
| **Treatmentsulpiride:Backtransfer** | 0.114 | 0.051 | 3560 | 2.218 | 0.027 |
| **Treatmentsulpiride:GenotypeA1-** | 0.03 | 0.072 | 72 | 0.42 | 0.676 |
| **Backtransfer:GenotypeA1-** | 0 | 0.051 | 3560 | -0.004 | 0.997 |
| **Treatmentsulpiride:Trustee** | -0.078 | 0.102 | 3560 | -0.759 | 0.448 |
| **Backtransfer:Trustee** | -0.027 | 0.076 | 3560 | -0.352 | 0.725 |
| **GenotypeA1-:Trustee** | -0.012 | 0.103 | 3560 | -0.112 | 0.911 |
| **Treatmentsulpiride:Backtransfer:GenotypeA1-** | -0.171 | 0.072 | 3560 | -2.363 | 0.018 |
| **Treatmentsulpiride:Backtransfer:Trustee** | -0.084 | 0.102 | 3560 | -0.819 | 0.413 |
| **Treatmentsulpiride:GenotypeA1-:Trustee** | 0.145 | 0.145 | 3560 | 1.003 | 0.316 |
| **Backtransfer:GenotypeA1-:Trustee** | 0.094 | 0.103 | 3560 | 0.917 | 0.359 |
| **Treatmentsulpiride:Backtransfer:GenotypeA1-:Trustee** | -0.002 | 0.145 | 3560 | -0.013 | 0.99 |

| ReciprocalTrial ~Treatment*Genotype*Trustee_c + (Trustee \|ID), family = binomial | Estimate | Est.Error | Q2.5 | Q97.5 |
| --- | --- | --- | --- | --- |
| **Intercept** | 0.634 | 0.133 | 0.377 | 0.894 |
| **Treatmentsulpiride** | 0.399 | 0.179 | 0.045 | 0.755 |
| **GenotypeA1M** | -0.015 | 0.179 | -0.37 | 0.335 |
| **Trustee** | -0.027 | 0.23 | -0.474 | 0.427 |
| **Treatmentsulpiride:GenotypeA1M** | -0.223 | 0.256 | -0.727 | 0.276 |
| **Treatmentsulpiride:Trustee** | -0.023 | 0.315 | -0.637 | 0.591 |
| **GenotypeA1M:Trustee** | 0.375 | 0.307 | -0.219 | 0.975 |
| **Treatmentsulpiride:GenotypeA1M:Trustee** | -0.103 | 0.439 | -0.965 | 0.76 |

Supplementary Table 5: Bayesian mixed logistic model predicting Reciprocal trials from drug treatment, genotype and trustee as predictors.

| ReciprocalTrial ~Treatment*Genotype*Trustee_c + (Trustee_c\|ID), family = binomial | Estimate | Std. Error | z value | Pr(>\|z\|) |
| --- | --- | --- | --- | --- |
| **(Intercept)** | 0.604 | 0.126 | 4.804 | <10e3 |
| **Treatmentsulpiride** | 0.392 | 0.171 | 2.296 | 0.022 |
| **GenotypeA1-** | -0.018 | 0.169 | -0.106 | 0.916 |
| **Trustee** | -0.049 | 0.22 | -0.222 | 0.824 |
| **Treatmentsulpiride:GenotypeA1-** | -0.217 | 0.24 | -0.903 | 0.366 |
| **Treatmentsulpiride:Trustee** | -0.009 | 0.299 | -0.03 | 0.976 |
| **GenotypeA1-:Trustee** | 0.38 | 0.295 | 1.289 | 0.197 |
| **Treatmentsulpiride:GenotypeA1-:Trustee** | -0.108 | 0.421 | -0.257 | 0.797 |

Supplementary Table 6: Frequentist model mixed logistic model predicting Reciprocal trials from drug treatment, genotype and trustee as predictors.

| ReciprocalTrial ~log(Serum)*Genotype*Trustee_c + (Trustee \|ID), family = binomial, data = Sulpiride group only | Estimate | Est.Error | Q2.5 | Q97.5 |
| --- | --- | --- | --- | --- |
| **Intercept** | 1.012 | 0.116 | 0.783 | 1.236 |
| **logserum_s** | **0.186** | **0.113** | **-0.04** | **0.41** |
| **GenotypeA1M** | -0.235 | 0.17 | -0.568 | 0.103 |
| **Trustee** | -0.058 | 0.175 | -0.404 | 0.282 |
| **logserum_s:GenotypeA1M** | -0.222 | 0.175 | -0.565 | 0.126 |
| **logserum_s:Trustee** | 0.325 | 0.169 | -0.003 | 0.661 |
| **GenotypeA1M:Trustee** | 0.263 | 0.256 | -0.241 | 0.774 |
| **logserum_s:GenotypeA1M:Trustee** | -0.429 | 0.256 | -0.935 | 0.074 |
| Supplementary Table 7: Bayesian mixed logistic model predicting Reciprocal trials Sulpiride group only, from serum levels (log-scaled), genotype and trustee as predictors.  MistakeTrial ~Logserum*Genotype*Trustee_c + (Trustee \|ID), family = binomial, data = Sulpiride group only | Estimate | Est.Error | Q2.5 | Q97.5 |
| **Intercept** | -2.068 | 0.248 | -2.589 | -1.607 |
| **logserum_s** | -0.331 | 0.242 | -0.818 | 0.143 |
| **GenotypeA1M** | 0.427 | 0.361 | -0.314 | 1.136 |
| **Trustee** | -0.156 | 0.233 | -0.622 | 0.300 |
| **logserum_s:GenotypeA1M** | **0.941** | **0.356** | **0.25** | **1.663** |
| **logserum_s:Trustee** | -0.57 | 0.231 | -1.048 | -0.127 |
| **GenotypeA1M:Trustee** | **-0.191** | **0.314** | **-0.812** | **0.428** |
| **logserum_s:GenotypeA1M:Trustee** | 0.595 | 0.315 | -0.01 | 1.227 |

Supplementary Table 8: Bayesian mixed logistic model predicting Mistake trials Sulpiride group only, from serum levels (log-scaled), genotype and trustee as predictors.

| Punishment ~ Treatment*Genotype + (1\|ID) | Estimate | Est.Error | Q2.5 | Q97.5 |
| --- | --- | --- | --- | --- |
| **Intercept** | 6.16 | 1.177 | 3.893 | 8.447 |
| **Treatmentsulpride** | 1.776 | 1.446 | -1.097 | 4.619 |
| **genotypeA1M** | -0.764 | 1.435 | -3.585 | 2.086 |
| **Treatmentsulpride:genotypeA1M** | -0.476 | 1.83 | -4.091 | 3.077 |

Supplementary Table 9: Negative reciprocity task: Bayesian mixed model predicting Punishment from treatment, genotype and trustee as predictors.

| Back-transfer ~ Treatment*Genotype + (1\|ID) | Estimate | Est.Error | Q2.5 | Q97.5 |
| --- | --- | --- | --- | --- |
| **Intercept** | 380.565 | 14.042 | 349.795 | 403.707 |
| **Treatmentsulpride** | -0.016 | 2.923 | -5.911 | 5.634 |
| **genotypeA1M** | -0.219 | 2.978 | -5.983 | 5.698 |
| **Treatmentsulpride:genotypeA1M** | -0.095 | 3 | -5.999 | 5.792 |

Supplementary Table 10: Positive reciprocity task: Bayesian mixed model predicting Back-transfer from treatment, genotype and trustee as predictors.

| PrecisionWeights_s ~ logserum_s * Genotype + (1 \| ID) | Estimate | Est.Error | Q2.5 | Q97.5 |
| --- | --- | --- | --- | --- |
| **Intercept** | 0.489 | 0.137 | 0.219 | 0.762 |
| **logserum_s** | **0.286** | **0.126** | **0.037** | **0.539** |
| **GenotypeA1M** | -1.118 | 0.2 | -1.507 | -0.721 |
| **logserum_s:GenotypeA1M** | -0.218 | 0.193 | -0.595 | 0.158 |

Supplementary Table 11: Bayesian mixed model predicting PrecisionWeights from log serum levels and genotype in the Sulpiride group only.

| prec_weights_s ~ logserum_s * Genotype + (1 \| ID) | Value | Std.Error | DF | t-value | p-value |
| --- | --- | --- | --- | --- | --- |
| **(Intercept)** | 0.511 | 0.13 | 1862 | 3.94 | <10e3 |
| **logserum_s** | 0.293 | 0.126 | 34 | 2.317 | 0.027 |
| **GenotypeA1-** | -1.165 | 0.194 | 34 | -6.005 | <10e3 |
| **logserum_s:GenotypeA1-** | -0.234 | 0.196 | 34 | -1.196 | 0.24 |

Supplementary Table 12: Frequentist mixed model predicting precision weights from log serum levels and genotype in the Sulpiride group only.

### Side Effects

The effect of sulpiride on blood-pressure and heart-rate, self-reported side-effects and mood were adopted from previous published work with the same cohort (Eisenegger et al., 2014; Naef et al., 2017). Supplementary Table 13 shows all side-effects measures, their changes over time, as well as the results of a Mann-Whitney test for differences across treatment groups. Significance levels are not above chance level if corrected for multiple testing (Holm-Bonferroni correction).

| side effects | time point | N | Plac. | Sulp. | Sign. (*p*) |
| --- | --- | --- | --- | --- | --- |
| Heart rate | base | 76 | 69.2 | 67.5 | 0.807 |
|  | 3 h | 76 | 63.8 | 64.9 | 0.596 |
|  | δ | 76 | -5.4 | -2.6 | 0.666 |
| Blood pressure systolic [mm hg] | base | 76 | 132.2 | 132.8 | 0.783 |
|  | 3 h | 76 | 128.1 | 127.5 | 0.975 |
|  | δ | 76 | -4.1 | -5.4 | 0.621 |
| Blood pressure diastolic [mm hg] | base | 76 | 76.1 | 76.9 | 0.856 |
|  | 3 h | 76 | 72.0 | 70.9 | 0.629 |
|  | δ | 76 | -4.1 | -6.0 | 0.240 |
| VAS: alertness (mean) | base | 76 | 22.6 | 23.4 | 0.880 |
|  | 3 h | 75 | 28.5 | 28.8 | 0.945 |
|  | δ | 75 | 5.9 | 5.7 | 0.719 |
| VAS: contentedness (mean) | base | 76 | 18.7 | 19.6 | 0.767 |
|  | 3 h | 75 | 20.6 | 22.1 | 0.660 |
|  | δ | 75 | 2.0 | 3.0 | 0.304 |
| VAS: calmness (mean) | base | 76 | 22.6 | 24.9 | 0.659 |
|  | 3 h | 75 | 23.0 | 23.4 | 0.812 |
|  | δ | 75 | 0.4 | -0.8 | 0.890 |
| NVL: any effect | 3h | 75 | -31.5 | -38.3 | 0.743 |
| NVL: bad effects | 3h | 75 | -42.4 | -43.1 | 0.439 |
| NVL: good effects | 3h | 75 | -40.2 | -40.7 | 0.570 |
| NVL: high | 3h | 75 | -43.3 | -41.6 | 0.204 |
| NVL: rush | 3h | 75 | -41.3 | -43.6 | 0.270 |
| NVL: like drug | 3h | 75 | -16.8 | -14.6 | 0.417 |
| NVL: stimulated | 3h | 75 | -39.4 | -36.7 | 0.100 |
| NVL: performance impaired | 3h | 75 | -38.2 | -35.6 | 0.152 |
| NVL: performance improved | 3h | 75 | -38.1 | -41.1 | 0.629 |
| NVL: willing to take again | 3h | 75 | 8.8 | 2.9 | 0.656 |
| NVL: willing to pay for | 3h | 75 | -39.3 | -40.8 | 0.402 |
| NVL: active-alert-energetic | 3h | 75 | -35.1 | -37.9 | 0.844 |
| NVL: shaky/jittery | 3h | 75 | -42.0 | -36.0 | 0.337 |
| NVL: euphoric | 3h | 75 | -43.3 | -38.7 | 0.158 |
| NVL: irregular or racing heart | 3h | 75 | -45.8 | -44.7 | 0.153 |
| NVL: talkative-friendly | 3h | 75 | -39.1 | -31.9 | 0.049 |
| NVL: nauseated, queasy or sick to stomach | 3h | 75 | -46.4 | -46.5 | 0.393 |
| NVL: nervous or anxious | 3h | 75 | -42.9 | -45.1 | 0.734 |
| NVL: restless | 3h | 75 | -36.2 | -30.8 | 0.085 |
| NVL: sluggish-lazy-fatigued | 3h | 75 | -25.7 | -23.6 | 0.487 |

Supplementary Table 13: Physiological and self-reported side effects following drug. (Notes. Base = baseline; 3h = 3 hours after drug loading; δ = difference between the value 3 hours after drug loading and the baseline; N = number of observations; Plac. = Placebo group; Sulp. = Sulpiride group; Sign. = Significance of Mann-Whitney tests for differences.)

### Supplementary Notes

**Supplementary Note 1**: Definition of reciprocal and mistake trials

$$Reciprocal Trial = ((Backtransfer==1 \& Change > 0) | (Backtransfer == 1 \& Investment == 10)) | ((Backtransfer==-1 \& Change < 0) | (Backtransfer == -1 \& Investment == 0)),$$

$$Mistake Trial = ((Backtransfer==1 \& Change < 0) | (Backtransfer == -1 \& Change > 0))$$

**Supplementary Note 2**: Outcome maximising agent

A rational agent, that does not take the uncertainty about their estimate into consideration, would at each trial choose the investment that brings the highest expected value. When the trustee betrays after an investment $I$, the participants ends up with an outcome of $10-I$. When the trustee equalizes both players receive half of the overall gain. The outcome $V$ in that case is:

$$V=\frac{I-10+10+ 3*I}{2}=10+I$$

If an agent has a subjective belief that the trustee will equalize with probability $p$, then the expected values of his investment $I$ is simply:

$$EV\left( I \right)=p*\left( 10+I \right)+\left( 1-p \right)*\left( 10-I \right)=10+I(2p-1)$$

Which is a linear function of $p.$ Meaning that as soon as $p$ is believed to be above $0.5$ the outcome maximising agent should give a $10$ and as soon as $p$ is below $0.5$ they should invest $0.$

### Supplementary Figures

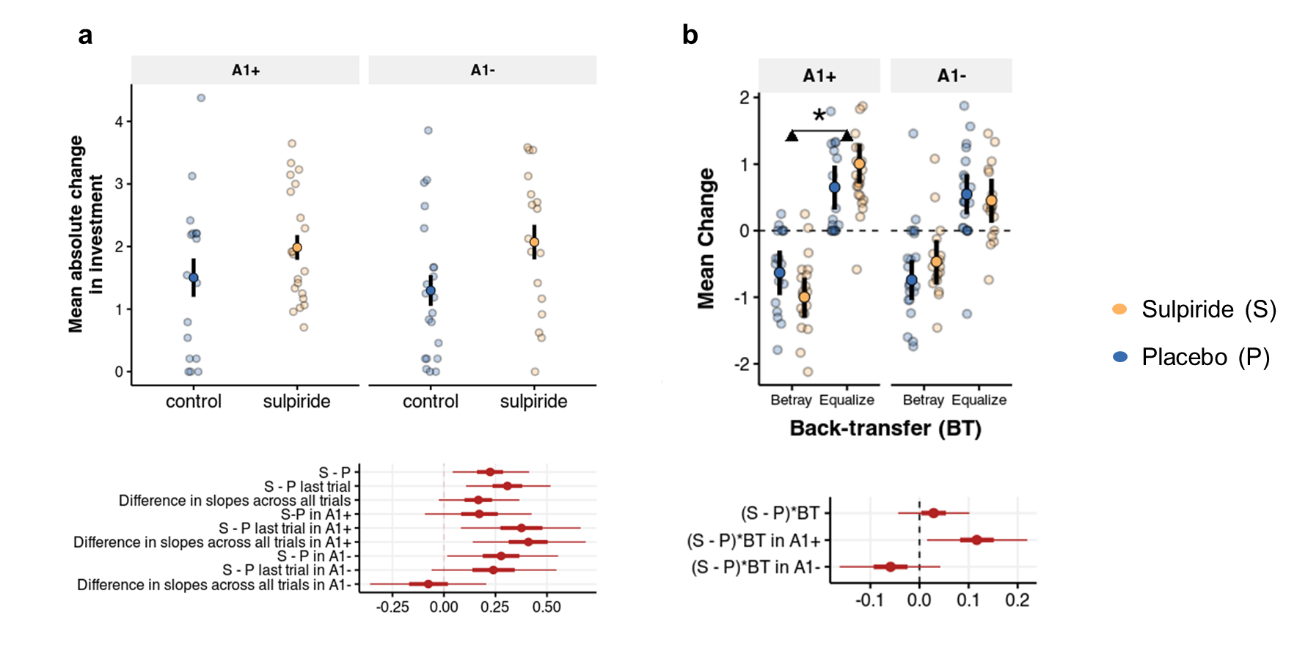

**Supplementary Fig. 1 | Effects of sulpiride on behaviour in the repeated trust game. a**, Mean and standard errors of absolute change of investment from one trial to the next for both treatment and genotype groups. 95% CrI of effect sizes show a main effect of sulpiride, no interaction effect of treatment and genotype, but a differential effect of sulpiride on the slope of two genotype groups. **b**, Mean and 95% CrI of relative change in investment following positive and negative feedback, for both genotype groups separately, predicted from a Bayesian multilevel model including Genotype as a predictor. Dots are raw means for each participant. Effect sizes below. The 95% CrI of the interaction effect of Drug and Back-transfer was above zero only in the A1+ group. We found no interaction effect of sulpiride and Back-transfer (b = 0.089, 95% CrI [-0.135, 0.314], P(b<0) = 0.217, d = 0.029, 95% CrI [-0.044, 0.102]), but observed a three-way interaction effect, with the Genotype variable (b = -0.545, 95% CrI [-0.987, -0.103], P(b>0) = 0.007, d = -0.176, 95% CrI [-0.319, -0.034]). In the A1+ group, participants administered the D2 antagonist tended to increase their investment following positive, and decrease their investment following negative, responses from the trustees (b = 0.361, 95% CrI [0.048, 0.68], P(b<0) = 0.013, d = 0.116, 95% CrI [0.016, 0.219]). There was no difference between sulpiride and placebo administration in A2 homozygotes (b = -0.179, 95% CrI [-0.502, 0.131] , P(b>0) = 0.127, d = -0.057, 95% CrI [-0.159, 0.042]).

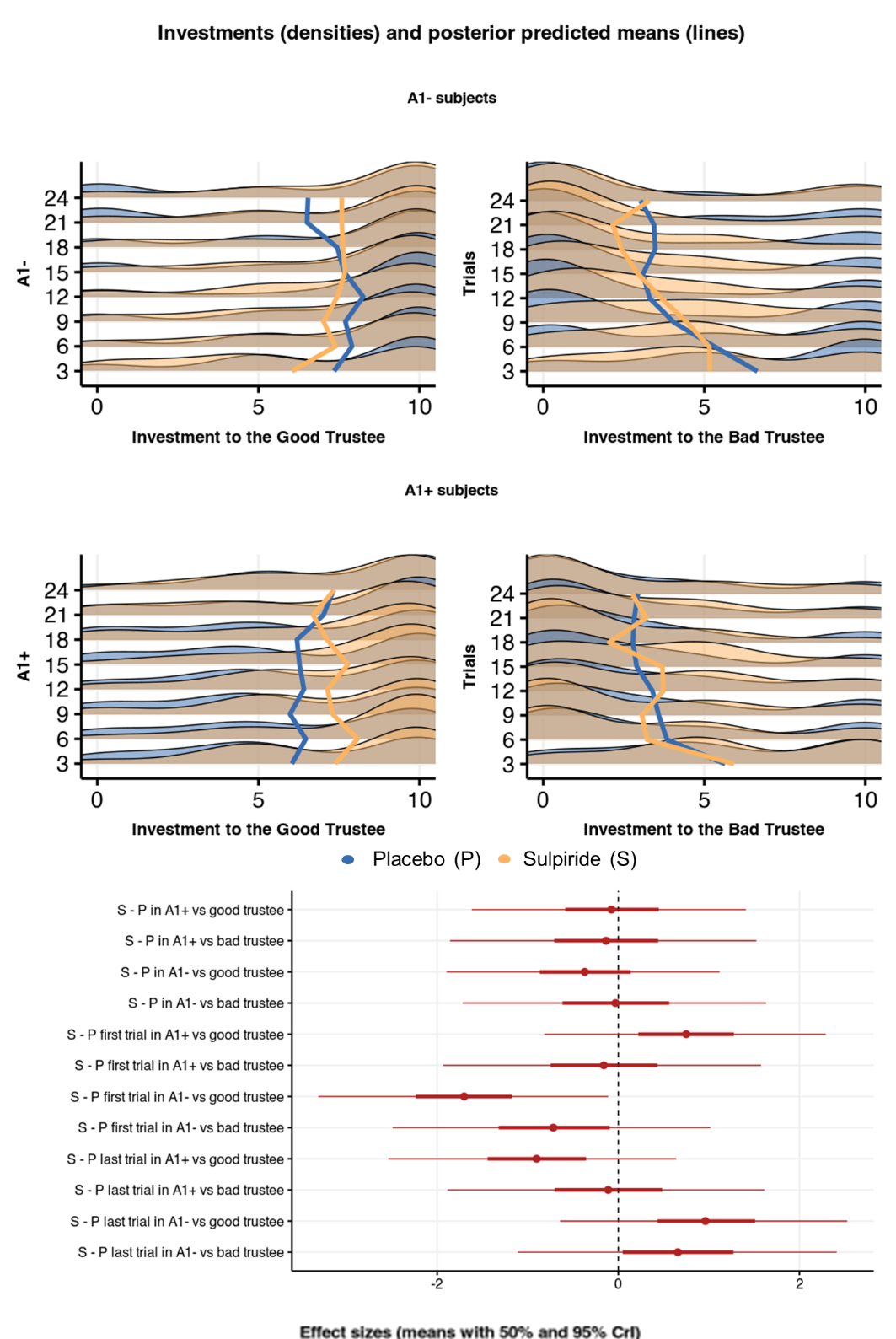

**Supplementary Fig. 2 | Effects of sulpiride on average investments in the repeated trust game.** Density plots of investments for placebo and sulpiride group overlayed and grouped within the trustee and genotype (binned across 3 trials for clarity). Lines are mean investments predicted from a multilevel Bayesian model that included Trial, Genotype, Trustee and Treatment as fixed and Trustee as participant-level random effects. Effect sizes are show below for average investments, as well as for differences in the first trial (initial trustworthiness) and in the last (learned trustworthiness).

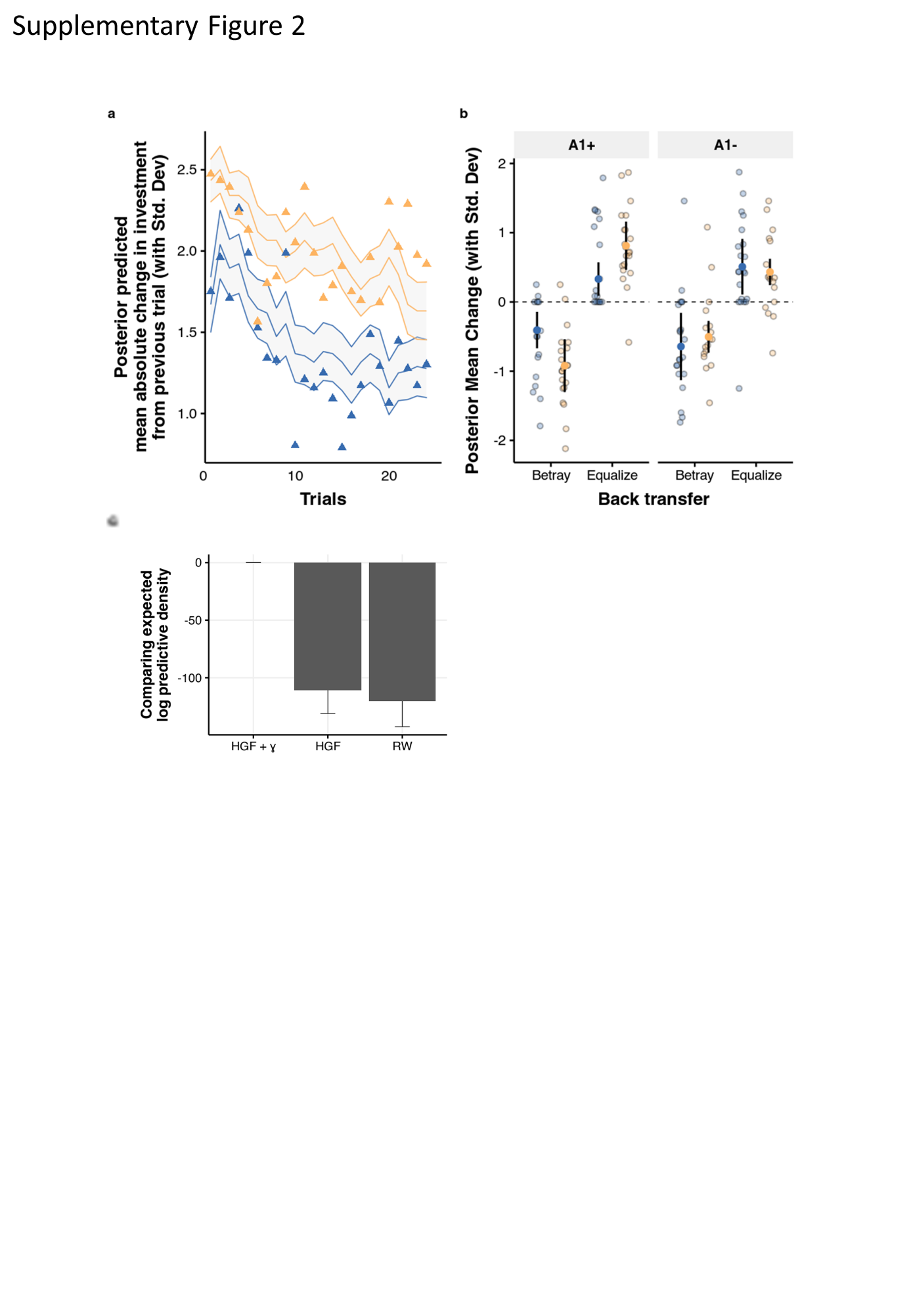

**Supplementary Fig. 3 | Model Validation and Model comparison. a,** Posterior predictive checks for absolute change of investment from one trial to the next, plotted over raw means. **b,** Posterior predictive check for the relative change in investment following positive or negative feedback plotted over raw means. **c,** Comparing the HGF model with all four parameters ($\omega$, $\mu_{0}$, $\eta$, $\gamma$) with an HGF model without $\gamma$ and a RW model. On the y-axis is the relative difference in the predictive density (on a log scale).

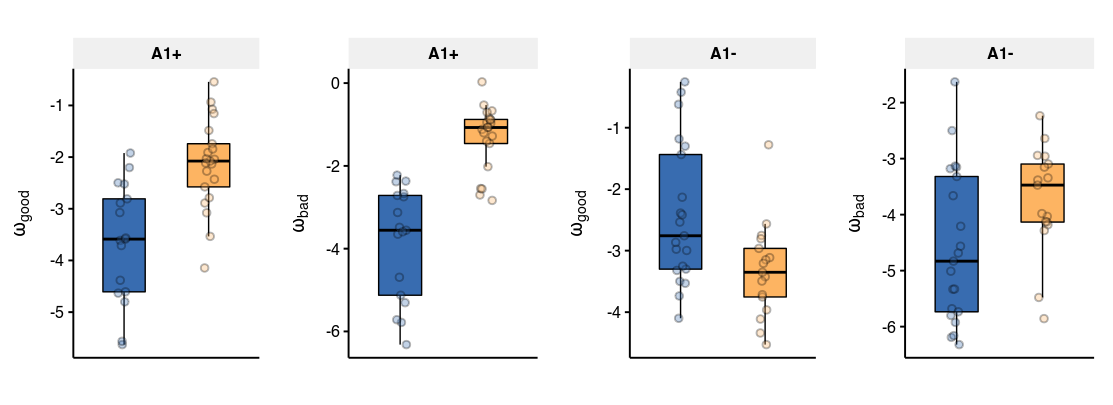

**Supplementary Fig. 4 | Comparing the effects of sulpiride on belief volatility across the two trustees**
